# Age-dependent spatial transfer of cortical harmonics to scalp EEG in infancy

**DOI:** 10.64898/2026.04.30.722070

**Authors:** Hyung G. Park

## Abstract

Infancy is a period of rapid change in cortical geometry and head volume conduction, raising a central question for developmental neuroimaging: how should spatial brain coordinates be defined so that they remain interpretable across age? In scalp electrophysiology, this question has two coupled aspects. First, age-dependent forward propagation through the head determines which cortical spatial patterns are expressed at the scalp. Second, cortical Laplace–Beltrami (LB) eigenmodes computed independently on age-specific templates need not remain directly comparable by mode index, even when they represent related low-dimensional structure. Using age-specific infant anatomical templates distributed through MNE-Python, we computed age-specific cortical LB eigenmodes, age-specific EEG forward models, and the corresponding forward-projected cortical harmonic dictionaries. We then quantified (i) age-dependent forward transfer to the scalp, (ii) cross-age mismatch in cortical basis, coefficient, and sensor spaces, and (iii) the improvement obtained from a simple local tracking procedure. Forward-projected scalp gain was consistently concentrated in lower cortical harmonic orders across infancy, with a broadly stable coarse-to-fine profile across age, although the absolute gain varies over age. In cortical mode space, independently computed neighboring-age cortical LB bases were only locally comparable by nominal mode index, exhibiting local reordering, mode-index drift, and coefficient leakage into nearby modes. This non-equivalence had practical consequences: neighboring-age infant dictionaries were not fully interchangeable across age in low-dimensional sensor space, and a fixed adult-derived basis was systematically suboptimal for recovering infant same-index harmonic coordinates, with the largest penalties in early-to mid-infancy. Sequential Procrustes tracking substantially improved same-index neighboring-age consistency. Together, these results motivate age-parameterized or explicitly tracked cortical harmonic coordinates for longitudinal developmental EEG and, more broadly, for lifespan analyses that seek geometrically meaningful spatial coordinates across changing brains.

## 1 Introduction

A central challenge in developmental neuroimaging is that the brain’s spatial coordinate system is itself developmentally dynamic. Across infancy and early childhood, cortical folding, cortical surface growth, head size, and scalp–brain geometry all change rapidly (Li et al., 2014; Beauchamp et al., 2011; O’Reilly et al., 2021). As a result, spatial patterns that are naturally defined on the cortical surface at one age need not be represented, observed, or interpreted in the same way at another age (O’Reilly et al., 2021; Lew et al., 2013). This issue is especially important for scalp electrophysiology, where measured potentials reflect not only cortical spatial patterning but also propagation through an age-dependent head volume conductor (Srinivasan et al., 1996, 1998; Lew et al., 2013). For longitudinal developmental studies, the problem is therefore not only how to represent cortical spatial structure at a given age, but also how to define spatial coordinates that remain interpretable across age (Hervé et al., 2022).

This problem is naturally connected to cortical harmonic coordinates. Cortical Laplace–Beltrami (LB) eigenmodes provide an intrinsic multiscale basis on the cortical surface, ordered from coarse to fine spatial structure (Reuter et al., 2006; Shi et al., 2009). In mature-brain settings, such eigenmodes have become increasingly useful as geometry-aware coordinates for describing large-scale brain activity and cortical spatial organization (Müller et al., 2022; Pang et al., 2023). More recently, forward-projected cortical eigenmodes have also been shown to provide an efficient and interpretable sensor-space representation of resting-state EEG in adults (Park, 2026). In developmental settings, however, two additional questions arise. First, because scalp EEG is spatially low-pass filtered by the head, which cortical harmonic orders are preferentially expressed at the scalp at each age? Second, even if age-specific cortical LB modes are well defined within each template, can their nominal indices be interpreted consistently across age, or does developmental geometric change induce local reordering and mixing that makes naive same-index comparison misleading?

Existing spatial-spectral work has shown that scalp EEG is dominated by low spatial frequencies and that volume conduction strongly limits recoverable fine-scale structure (Srinivasan et al., 1996, 1998; Nunez and Srinivasan, 2006). Recent work has extended such analyses to realistically shaped head models using SPHARA and BEM, further underscoring how head geometry constrains scalp spatial structure (Ahlfors et al., 2010; Graichen et al., 2025). At the same time, geometry-aware studies suggest that cortical eigenmodes provide a parsimonious scaffold for large-scale brain activity and cortical organization (Müller et al., 2022; Pang et al., 2023; Park, 2026). What remains unresolved is how these ideas extend across rapidly changing infant anatomy. In particular, it is still unclear how strongly lower-order cortical harmonics dominate infant scalp observations, whether independently computed age-specific harmonic coordinates are interchangeable across age, and whether any mismatch is large enough to matter in practice for sensor-space representation and coordinate recovery.

In this paper, we address these questions in a template-based developmental framework. Using age-specific infant anatomical templates, we compute cortical LB eigenmodes, age-specific EEG forward models, and forward-projected cortical harmonic dictionaries (Richards et al., 2016; O’Reilly et al., 2021; Gramfort et al., 2014). We first characterize the forward transfer of age-specific cortical harmonics to the scalp. We then examine the practical consequences of developmental mismatch, including neighboring-age sensor-space non-equivalence and the performance of a fixed adult-derived basis for recovering infant same-index harmonic coordinates. Finally, we study cross-age comparability in cortical basis, coefficient, and sensor spaces, and evaluate a simple local tracking procedure to improve cross-age consistency. Our goal is to determine whether age-specific cortical harmonic coordinates can serve as a stable developmental representation, or whether longitudinal developmental EEG requires age-parameterized or explicitly tracked harmonic coordinates.

## 2 Data and anatomical templates

Our primary data source consists of the age-specific infant templates distributed through MNE-Python, spanning 13 ages from 2 weeks to 2 years (`2wk, 1mo, 2mo, 3mo, 4.5mo, 6mo, 7.5mo, 9mo, 10.5mo, 12mo, 15mo, 18mo, 2yr`) (Richards et al., 2016; O’Reilly et al., 2021; Gramfort et al., 2014). Each template provides age-specific cortical surfaces, a source space, and a three-layer BEM solution, enabling a fully age-specific forward-modeling pipeline within a common software framework.

For the primary analyses, we use the template white surfaces as the cortical manifolds on which the intrinsic LB eigenmodes are defined, and the template `oct6` source spaces to construct age-specific forward-projected dictionaries. The full cortical surface is the natural object on which the intrinsic LB basis is defined, whereas the source space is the discrete set of vertices on which the EEG forward operator is evaluated. To ensure exact compatibility with the leadfield matrix, we restrict the full-surface eigenmodes to the source-space vertices used by the age-specific forward model. For separate cortical correspondence analyses, however, we also compute eigensystems on the full cortical mesh rather than only on the source-space subsampling, so that cross-age basis mismatch is not confounded with template-specific differences in source-space discretization or age-varying source pruning.

Our primary sensor-space analysis uses the built-in template 10–20 montage distributed with each infant template, yielding *M* = 20 EEG channels in age-specific template coordinates (O’Reilly et al., 2021; Jurcak et al., 2007). Although sparse, this age-appropriate montage provides a controlled setting for evaluating forward transfer, developmental basis mismatch, and neighboring-age sensor-space non-equivalence. Denser infant montages would be valuable, but age-specific dense sensor configurations are not currently distributed as part of the template package, so we leave that extension for future work.

## 3 Methods

Our analyses address three linked questions: how age-specific cortical harmonics are transferred to the scalp, how developmental mismatch affects practical sensor-space representation and recovery, and how comparable independently computed age-specific harmonic coordinates remain across infancy. To address these questions, we compute age-specific cortical LB eigenmodes and forward-projected cortical harmonic dictionaries, quantify forward-projected scalp gain across harmonic order, and evaluate developmental mismatch in sensor, basis, and coefficient spaces. We then examine whether cross-age instability is concentrated in locally crowded eigenvalue neighborhoods, distinguish cortical-basis and forward-operator-side contributions to neighboring-age sensor mismatch, and assess Sequential Procrustes tracking as a local remedy for improving same-index neighboring-age consistency. Together, these analyses distinguish forward observability, practical recoverability, and developmental coordinate comparability.

### 3.1 Age-specific cortical geometric eigenmodes and their physical scale

For each template age *a*, we compute cortical Laplace–Beltrami (LB) eigenmodes 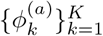 on the age-specific cortical surface, using *K* = 50 modes per hemisphere. These modes provide an intrinsic multiscale coordinate system on the cortex, ordered from coarse to fine spatial variation. We write the resulting age-specific cortical harmonic basis as

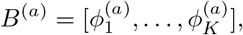

where 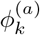 is the *k*-th cortical geometric eigenmode and 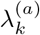 is the corresponding eigenvalue. The modes are obtained by solving

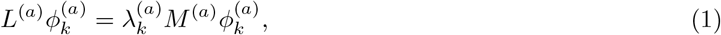

where *L*^(*a*)^ is the cotangent Laplacian and *M* ^(*a*)^ is the area-weighting mass matrix on the age-*a* cortical mesh. The eigenvectors are normalized with respect to the area-weighted inner product. Implementation details for mesh construction, discrete operators, normalization, and source-space restriction are given in Supplementary Section S1.

Because the eigenvalues 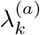 increase with spatial complexity, the mode index *k* can be interpreted as a harmonic order, with lower *k* corresponding to spatially smoother cortical patterns and higher *k* corresponding to finer spatial variation. To summarize how the physical meaning of a fixed nominal mode order changes across development, we define the wavelength-like scale

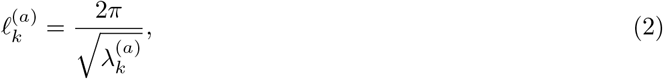

which serves as an intuitive proxy for the physical spatial scale associated with mode *k* on the age-*a* cortex: larger values of 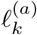 correspond to coarser cortical patterns and smaller values to finer ones. Because overall cortical size changes substantially across age, we also consider the area-normalized quantity

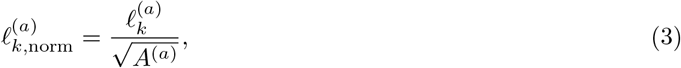

where *A*^(*a*)^ is the hemisphere-specific cortical surface area at age *a*. These quantities are used in Section 4.1 to assess how the physical interpretation of a fixed nominal mode order changes across development and to distinguish global size effects from residual departures from exact same-index comparability.

### 3.2 Forward projection to scalp and scalp-gain summaries

For each age *a*, let 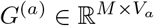 denote the fixed-orientation EEG leadfield matrix mapping source amplitudes at the active source-space vertices to the *M* = 20 scalp channels; the number of active source vertices, denoted *V*_*a*_, varies across age. The forward model is computed from the age-specific source space, age-specific three-layer BEM solution, and age-specific template 10–20 montage (Richards et al., 2016; O’Reilly et al., 2021; Gramfort et al., 2014).

For forward-model analyses, we restrict the full-mesh cortical harmonic basis *B*^(*a*)^ to the source-space vertices used by the age-specific leadfield. Denoting the restricted basis by 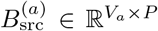, we define the age-specific forward-projected cortical harmonic dictionary as

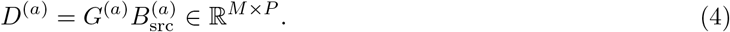

Each column of *D*^(*a*)^ is the scalp topography produced by one cortical harmonic under the age-specific forward model. In the current implementation, *P* = 2*K*, with *K* left-hemisphere columns followed by *K* right-hemisphere columns. After restriction to the source space, each basis column is re-normalized to unit Euclidean norm so that forward-projected column norms are directly comparable across age and are not driven by age-varying source-vertex count.

To summarize deterministic forward transfer to the scalp, we define the scalp gain of each forward-projected cortical harmonic by

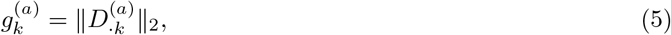

which directly measures how strongly the *k*-th cortical harmonic is expressed at the scalp at age *a*. For display, left-and right-hemisphere modes of the same nominal order are combined, and the resulting age-specific gain profile is normalized to sum to one across harmonic order. Here, nominal mode order refers to the mode index after sorting the eigenvalues from smallest to largest, so lower-order modes correspond to spatially coarser cortical patterns and higher-order modes to finer ones. We also compute cumulative summaries, including the smallest harmonic mode order accounting for 50%, 70%, and 90% of total forward-projected scalp gain. These summaries quantify whether scalp EEG primarily reflects a relatively small low-order portion of the cortical harmonic hierarchy or whether appreciable forward-projected gain extends to finer cortical spatial scales.

### 3.3 Developmental basis mismatch in sensor space

We study the practical sensor-space consequences of developmental basis mismatch in two related analyses.

First, as a neighboring-age benchmark, we assess whether neighboring-age infant dictionaries can be used interchangeably to represent low-dimensional age-specific scalp patterns. For an age-specific dictionary *D*^(*a*)^ from Equation (4), we generate low-dimensional sensor patterns

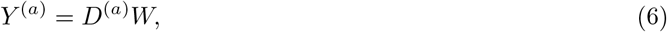

where *W* is a matrix of simulated coefficient vectors restricted to the leading 5 modes per hemisphere. The coefficients in *W* are generated as random draws with equal variance across the retained coordinates, yielding 1000 sensor patterns at each age. We then project these patterns onto the within-age sensor subspace, span(*D*^(*a*)^), and onto a neighboring-age sensor subspace, span(*D*^(*b*)^), and compare the resulting representation quality,

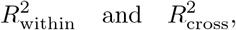

together with the loss

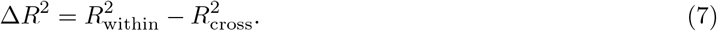

We also compute mean principal angles and projector distances between the corresponding sensor subspaces. Details of the *R*^2^ calculation and projector-distance summary are given in Supplementary Section S3.

Second, we assess whether a fixed adult-derived analysis basis can recover infant same-index harmonic coordinates as effectively as the correct age-specific infant basis. For each infant template age *a*, let *B*^(*a*)^ denote the age-specific infant cortical harmonic basis restricted to the age *a*-specific EEG source space, and let *G*^(*a*)^ denote the corresponding age-specific forward operator. The matched infant analysis dictionary is

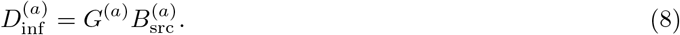

To study developmental basis mismatch, we also construct an adult-derived analysis basis on the same infant geometry. Starting from a standard adult cortical harmonic basis *B*_adult_, we map it to the age-*a* infant cortical mesh using the spherical-registration-based mapping, restrict it to the age-*a* source space, re-normalize its columns, and sign-align the mapped columns to the age-specific infant basis by nominal mode index. We denote the resulting adult-derived basis on infant age *a* by *B*_adult→*a*_, and define

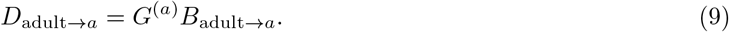

Because both 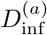 and *D*_adult→*a*_ use the same age-*a* forward operator, differences between them isolate cortical basis mismatch rather than forward-model mismatch. Details of this construction are given in Supplementary Section S5.

For each age *a*, we simulate latent infant-basis coefficient trajectories *w*^(*a*)^(*t*) and generate noisy scalp observations

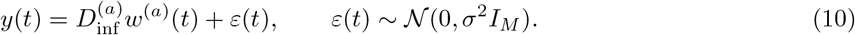

The coefficient trajectories *w*^(*a*)^(*t*) are generated as independent AR(1) processes with equal marginal variance over the first 10 modes per hemisphere. For the matched benchmark, coefficients are recovered using the age-specific infant dictionary 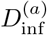. In contrast, for the adult-mismatch analysis, coefficients are recovered using the adult-derived dictionary *D*_adult→*a*_. In both cases, recovery is performed using the posterior mean under a Gaussian prior:

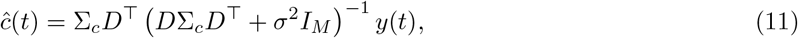

where *D* is the analysis dictionary and Σ_*c*_ is diagonal with the same variance structure used to generate the latent coefficients. Recovered coefficients are also compared directly with the true infant coefficients *w*^(*a*)^(*t*) by nominal mode index. This asks whether a fixed adult-derived analysis basis can recover infant harmonic coordinates of the same nominal order. We summarize recoverability using mode-wise correlation between true and recovered coefficients, the mean recoverability of the first 10 paired harmonic orders, and the correct-minus-adult-derived recoverability drop by age and harmonic mode order. In (10), the sensor-level signal to noise ratio (SNR) is defined in terms of the expected average signal power per channel; 0 dB corresponds to equal expected signal and noise power, 10 dB to tenfold higher signal power, and 20 dB to one-hundredfold higher signal power.

### 3.4 Cross-age cortical eigenmode correspondence

Independently computed age-specific cortical LB eigenmodes need not remain directly comparable by mode index, even when the underlying low-dimensional harmonic organization is broadly preserved. The main reasons are local rotations within nearly degenerate neighborhoods of nearby modes with similar eigenvalues and developmental geometric change that induces local reordering of nearby modes. Thus, comparing “mode *k*” at age *a* with “mode *k*” at age *b* is not guaranteed to be meaningful unless some form of age-parameterization or tracking is imposed.

To quantify this mismatch, we compute neighboring-age “correspondence” matrices in cortical mode space. Let *B*^(*a*)^ denote the full-mesh cortical basis at age *a*. We map the basis from age *b* onto the cortical mesh of age *a* using the template spherical registration (Dale et al., 1999; Fischl et al., 1999). Specifically, each template provides a spherical-registration surface that aligns cortical topology to a common spherical coordinate system; we transfer mode values from age *b* to the vertex locations of age *a* by nearest-neighbor interpolation on this registered sphere. Details of this mapping are given in Supplementary Section S2. Writing the mapped basis as *B*^(*b*→*a*)^, we define the cortical correspondence matrix

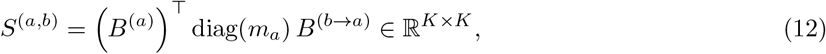

where *m*_*a*_ is the vector of vertex-area weights on age *a*, i.e., the diagonal entries of the mass matrix *M* ^(*a*)^ in (1).

Since the sign of each LB eigenfunction is arbitrary and does not carry cross-age interpretive meaning, we analyze the absolute correspondence matrix |*S*^(*a,b*)^|. For a given evaluation-age mode *k*, its *best match* is defined as

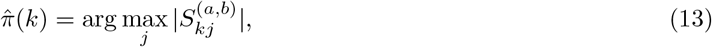

that is, the mode at age *b* (after mapping to the age-*a* mesh) whose absolute correspondence with mode *k* at age *a* is largest. Using (13), we summarize neighboring-age basis mismatch by: (i) the exact best-match rate, which is the proportion of modes for which 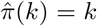 (ii) the near-neighbor match rates, which is the proportions with 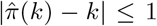 and 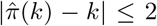 (iii) the mean absolute shift, which is the average of 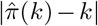 across modes; and (iv) adjacent reorderings, which count local departures from monotone ordering in the best-match sequence. Each neighboring-age pair is evaluated in both directions, and the directional summaries are then averaged.

We further assess how much the coordinates (i.e., coefficients) of the same cortical pattern change when re-expressed in a neighboring-age basis. We define the cross-age coefficient transfer operator

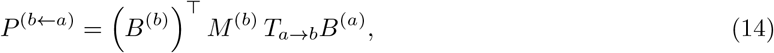

where *T*_*a*→*b*_ maps a cortical pattern from the age-*a* mesh to the age-*b* mesh via spherical-registration-based interpolation; details of the mapping operator *T*_*a*→*b*_, the role of the area-weight matrices, and the full coefficient-space formulation are given in Supplementary Section S4. If *c*^(*a*)^ ∈ ℝ^*K*^ denotes a coefficient vector at age *a*, then the corresponding coefficients at age *b* are

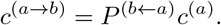

If *P* ^(*b*←*a*)^ were close to the identity, then same-index coefficients would remain stable across age; off-diagonal structure indicates coefficient mixing across nearby or more distant modes.

We evaluate two families of test patterns. First, *single-mode test patterns*, in which *c*^(*a*)^ = *e*_*k*_, isolate how an individual (i.e., the *k*th) nominal mode spreads across age. Second, *neighboring-mode test patterns*, in which the coefficient vector is concentrated in a small neighborhood around mode *k*, test whether a locally concentrated harmonic pattern remains concentrated nearby after age mismatch. For both families, we summarize and report: (1) the fraction of squared target coefficient mass remaining at the same nominal mode index; (2) the fractions remaining within *±*1 and *±*2 neighboring mode indices; (3) the peak shift, defined as the absolute displacement of the highest-mass target coefficient index from the nominal source index; and (4) the centroid shift, defined as the absolute displacement of the mass-weighted center of the target coefficient distribution from the nominal source index.

### 3.5 Local eigenvalue spacing (“mode crowding”) and cross-age instability

To understand why some cortical eigenmodes are harder to match across age than others, we quantify how crowded each mode is by nearby eigenmodes in the ordered LB spectrum. For mode *k* at age *a*, let 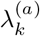 denote its eigenvalue and define the “local eigenvalue spacing”

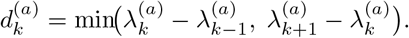

(For the first and last retained modes, we use the available one-sided spacing.) We then define the relative local spacing

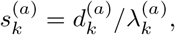

and the corresponding “crowding score”

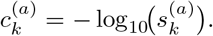

Smaller relative spacing indicates that a mode lies in a more crowded neighborhood of nearby eigenmodes with similar eigenvalues, so larger values of 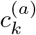 indicate greater crowding.

We then ask whether evaluation-age modes with larger crowding scores are less likely to preserve exact same-index correspondence across age. For each neighboring-age comparison—that is, evaluating on age *a* while comparing age-*a* modes to the age-*b* modes after mapping them onto the age-*a* mesh, or vice versa—we link the crowding score of each evaluation-age mode to its best-match index in the mapped neighboring-age basis. The best-match index for evaluation-age mode *k* is defined as (13), where *S*^(*a,b*)^ is the cross-age correspondence matrix defined in Equation (12). The primary summary outcome is exact-match probability as a function of crowding score, and we also report near-*±*1 and near-*±*2 match probabilities. These summaries are reported in Section 4.5.

### 3.6 Comparing cortical-basis and forward-operator contributions

To assess whether cross-age sensor-space mismatch is driven more strongly by changes in the cortical LB eigenmode basis or by changes on the forward-operator side, we performed one-component-at-a-time substitution comparisons. For age *a*, the native sensor dictionary is 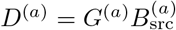 from Equation (4). The full neighboring-age comparison (between *a* and *b*) is therefore the comparison between the two native age-specific sensor dictionaries *D*^(*a*)^ and *D*^(*b*)^.

For each neighboring-age pair *a, b*, we constructed two partial substitutions. In the *cortical-basis substitution*, we replaced the age-*a* cortical basis by the age-*b* basis mapped to the age-*a* source space, while holding the age-*a* forward operator fixed:

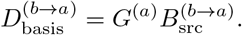

Here (*b* → *a*) indicates that the age-*b* cortical basis has been (mismatchingly) mapped onto the age-*a* source-space geometry before constructing the dictionary. In the *forward-operator substitution*, on the other hand, we held the age-*a* cortical basis fixed, mapped it to the age-*b* source space, and then applied (mismatchingly) the age-*b* forward operator:

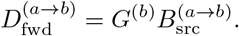

These analyses ask whether replacing one side of the age-specific dictionary, while holding the other fixed, produces a mismatch of similar magnitude to the full native neighboring-age dictionary comparison.

We quantify mismatch using the same low-dimensional simulated sensor patterns as in Section 3.3. For each comparison, we compute the representation loss 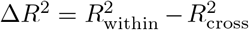 as in (7). We also compute the projector distance (using the Frobenius norm)

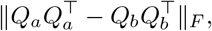

where *Q*_*a*_ and *Q*_*b*_ are orthonormal bases for the two sensor subspaces being compared.

### 3.7 Sequential Procrustes tracking of neighboring-age harmonic bases

As a simple local alignment procedure to improve cross-age comparability of age-specific cortical eigenmode coordinates, we applied *Sequential Procrustes tracking* to the age-specific cortical LB eigenmode bases. The goal of this procedure is not to define a final developmental coordinate system, but to provide a local alignment step that reduces neighboring-age mode reordering and rotation.

Let *a*_(1)_ *< a*_(2)_ *<* · · · *< a*_(*T*)_ denote the ordered template ages, from youngest to oldest, and let 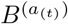 denote the corresponding cortical LB eigenmode basis at age *a*_(*t*)_. We aligned these bases sequentially, separately for each hemisphere, using the youngest age *a*_(1)_ as the initial reference. At each step, the native basis at age *a*_(*t*+1)_ was first mapped onto the cortical mesh of the previous age *a*_(*t*)_ using the template spherical registration and nearest-neighbor interpolation on the registered spherical surface (`sphere.reg`; see Dale et al., 1999; Fischl et al., 1999 and Supplementary Sections S2 and S6).

Because the observed cross-age mismatch was often local rather than global, we aligned the basis in small contiguous groups of nearby modes. In the current implementation, we used groups of three consecutive modes. Within each group, we solved a weighted orthogonal Procrustes problem on the previous-age mesh,

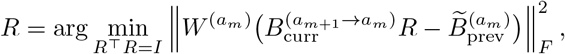

using the square-root area weights 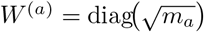, where *m*_*a*_ is the vertex-area weight vector introduced in Equation (12). The resulting orthogonal rotation *R* was then applied to the corresponding group of native modes at age *a*_(*t*+1)_, and the aligned columns were sign-adjusted so that, after mapping to the previous age, their area-weighted correspondence with the previously tracked columns was positive. Full implementation details are given in Supplementary Section S6.

This procedure yields a *Procrustes-tracked eigenmode basis* 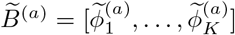 at each age. Because the procedure applies orthogonal rotations within local groups of nearby modes, the resulting columns are generally not exact Laplace–Beltrami eigenvectors. We therefore interpret 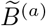 not as a strict tracked eigensystem, but as a locally aligned cortical eigenmode basis that preserves the relevant low-dimensional harmonic subspaces while improving cross-age coordinate consistency.

To evaluate whether Sequential Procrustes tracking improved neighboring-age comparability, we recomputed the same cross-age cortical correspondence summaries defined in Section 3.4 before and after tracking. For each neighboring-age pair and each directional evaluation, we computed the absolute correspondence matrix, exact same-index match rate, near-*±*1 and near-*±*2 match rates, and mean absolute best-match shift. Here, “directional” means that for a neighboring-age pair (*a, b*) we evaluate correspondence once on the age-*a* mesh after mapping age-*b* modes onto age *a*, and once on the age-*b* mesh after mapping age-*a* modes onto age *b*. Results were averaged across both directions and then across hemispheres for the main summaries. As a supplementary downstream validation, we also repeated the neighboring-age coefficient-space mismatch analysis using the tracked basis and compared exact same-index retained fraction before versus after tracking.

## 4 Results

### 4.1 Age-specific cortical harmonic coordinates co-move with cortical growth

Figures 1 and 2 provide complementary views of how cortical harmonic coordinates change across development. First, the modes visibly co-move with the age-specific cortical geometry: as cortical size and shape change, the harmonic patterns change with the surface rather than remaining tied to a fixed external coordinate system. Second, although the same nominal mode order is often broadly recognizable across age (across rows), its spatial pattern is not preserved exactly. Thus, even qualitatively, same nominal index modes are only approximately comparable across development.

**Figure 1:**
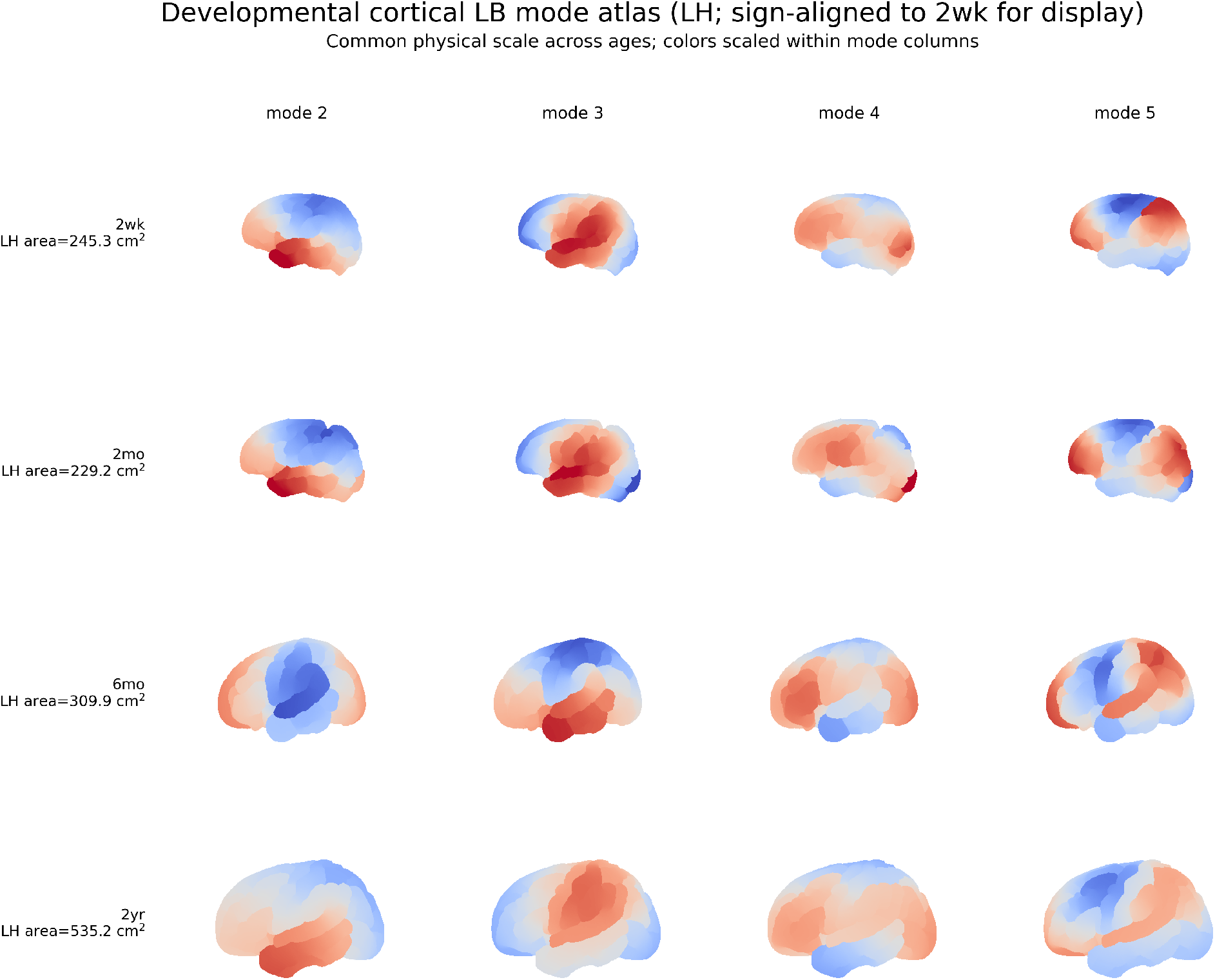
Developmental cortical LB atlas on age-specific geometry. Selected nominal cortical Laplace–Beltrami (LB) modes (*k* = 2, 3, 4, 5) are shown on age-specific left-hemisphere cortical surfaces at representative infant ages. LB eigenmodes were computed on the age-specific white surfaces and rendered on the corresponding pial surfaces for visualization. Within each mode column, the sign was aligned to the 2-week template using the registered spherical surfaces for display; this removes the arbitrary global sign ambiguity but does not enforce the requirement that mode *k* at one age should correspond to mode *k* at another age. All panels share a common physical scale, so cortical growth is visible across rows, and the row labels report left-hemisphere surface area. The figure shows that cortical harmonic coordinates co-move with the developing cortex geometry and therefore reflect an age-specific notion of spatial scale. At the same time, the spatial pattern associated with a fixed nominal mode order changes across age, illustrating why independently computed same-index modes are not automatically comparable without explicit age-parameterization or tracking.

**Figure 2:**
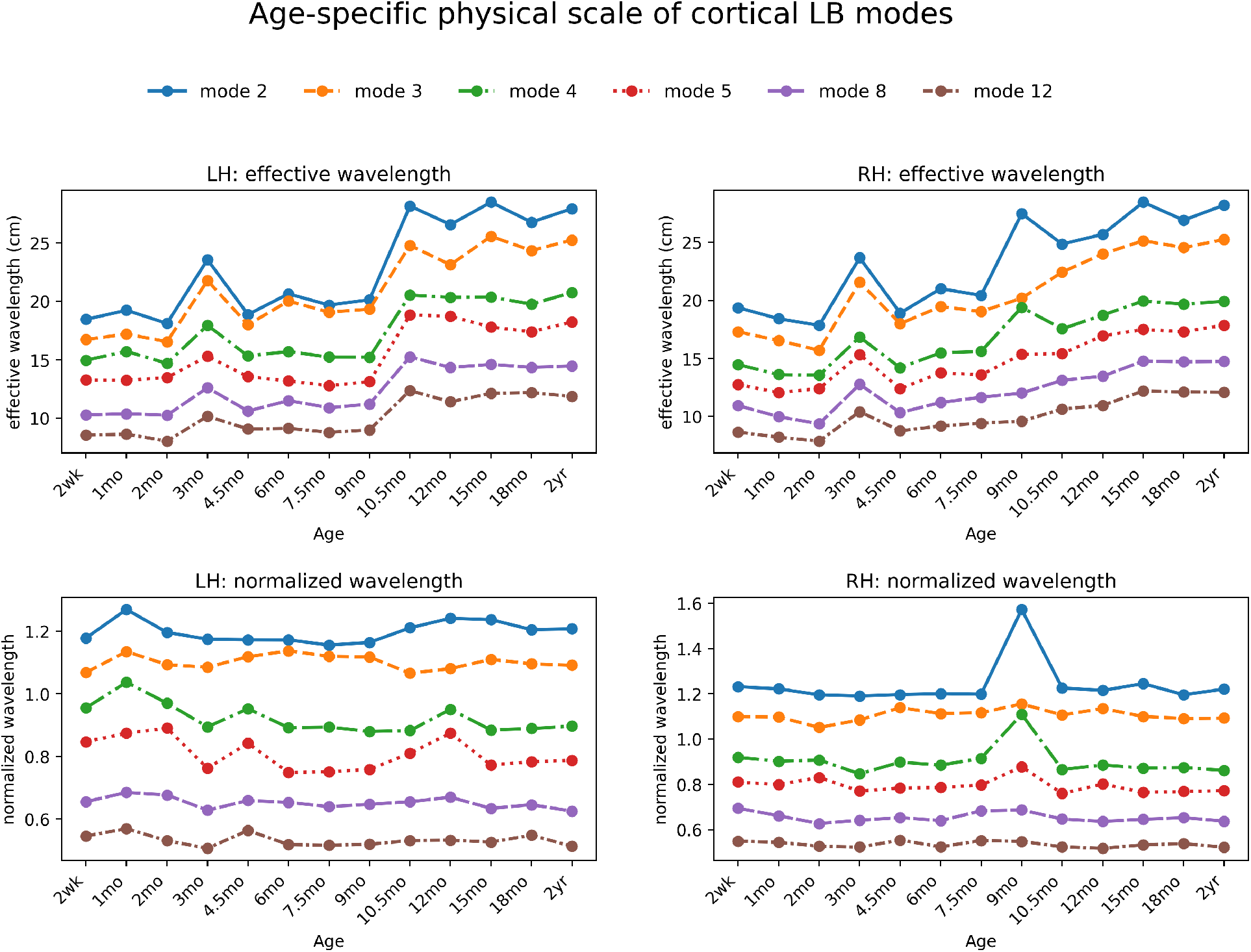
Age-specific physical scale of cortical LB modes. For selected nominal mode orders *k*, the top row shows the wavelength-like scale 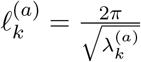, for the left and right hemispheres across age, where 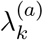 is the LB eigenvalue at age *a*. The bottom row shows the corresponding area-normalized quantity 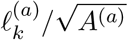, where *A*^(*a*)^ is the hemisphere-specific cortical surface area. The top row shows that, for a fixed nominal mode order, older brains tend to have larger wavelength-like scales, reflecting cortical growth. The substantially flatter trajectories after normalization (bottom row) indicate that much of this age dependence is driven by overall cortical size. Residual deviations from flatness, however, show that independently computed same-index modes are only approximately, not exactly, comparable across age, consistent with the cross-age correspondence analyses and motivating age-parameterized or explicitly tracked cortical harmonic coordinates.

Figure 2 quantifies this point more directly. For a fixed nominal mode order *k*, the corresponding wavelength-like scale 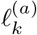 from Equation (2) increases with age, reflecting overall cortical growth. However, after normalization by 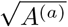 as in Equation (3), the trajectories become markedly flatter, indicating that much of this age dependence is driven by cortical size. The remaining deviations from flatness, however, show that independently computed same-index modes are only approximately, not exactly, comparable across age. These observations are consistent with the more formal cross-age cortical correspondence results reported in Section 4.4.

### 4.2 Forward-projected scalp gain is concentrated in lower cortical harmonic orders

Figure 3 summarizes, for each age (*a*), the distribution across cortical harmonic mode order (*k*) of the scalp-gain quantity 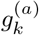 defined in Equation (5). This quantity measures how strongly a unit-norm cortical harmonic of order *k* is expressed at the scalp through the age-specific forward model and therefore provides a summary of which cortical spatial scales are preferentially transferred to scalp EEG. For display, left-and right-hemisphere modes of the same nominal order were combined, and the resulting age-specific gain profile was normalized to sum to one across the modes. This normalization emphasizes how scalp gain is distributed across harmonic order rather than absolute magnitude differences across ages.

**Figure 3:**
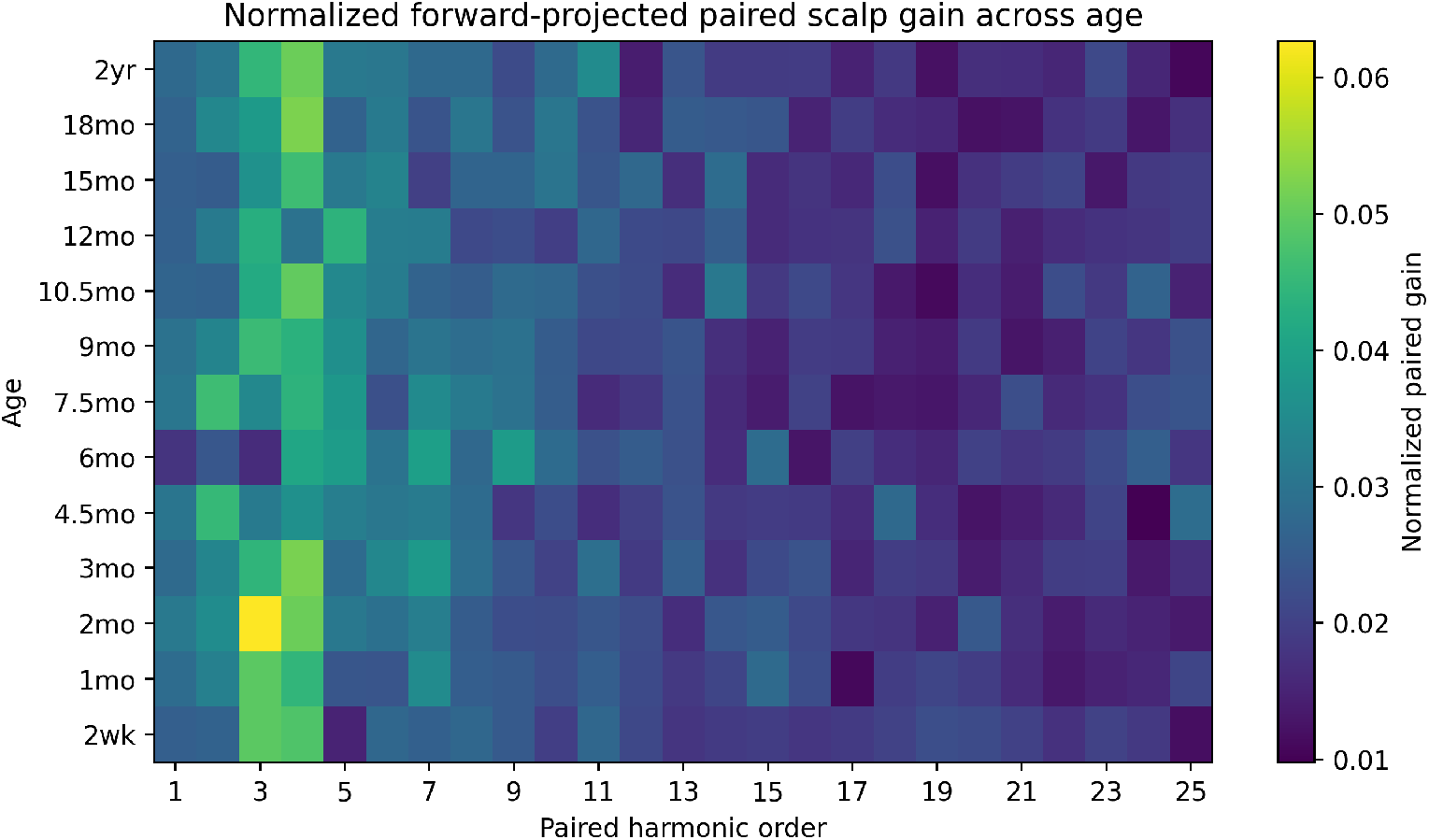
Normalized distribution of forward-projected scalp gain across age and cortical harmonic mode order. For each age, left-and right-hemisphere modes of the same nominal order were first combined, and the resulting gain profile was normalized to sum to one across harmonic order. The heatmap therefore shows how the age-specific scalp-gain quantity 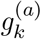 is distributed from lower to higher cortical harmonic orders, rather than absolute gain magnitude. Across infancy, the strongest forward-projected scalp gain is concentrated in lower harmonic orders, with substantially weaker gain at higher orders, indicating a broadly stable low-order-dominant forward-transfer profile across age. For visual clarity, the figure shows the first 25 harmonic orders after left-right pairing; the cumulative summaries reported in the text were computed using all 50 such orders.

Across infancy (2 weeks to 2 years), the strongest forward-projected gain is concentrated in lower cortical harmonic mode orders, with substantially weaker gain at higher mode orders. This low-order dominance is broadly stable across age, although the exact profile is not identical at every age. The corresponding cumulative summaries were also highly stable: approximately 18–21 harmonic orders (out of the total 50 modes) accounted for 50% of total forward-projected gain, 30–33 accounted for 70%, and 43–45 accounted for 90%. Thus, although absolute forward-projected gain may vary with age, the relative coarse-to-fine transfer profile remains consistently dominated by lower-order cortical harmonics throughout infancy. This pattern is consistent with the expected low-pass nature of scalp EEG. Although it does not by itself determine coefficient recoverability, it identifies which cortical spatial scales are preferentially transmitted to the scalp.

### 4.3 Developmental basis mismatch has practical consequences in sensor space

We next asked whether developmental changes in the cortical harmonic coordinate system have practically meaningful consequences in sensor space. We considered this question at two levels. First, as a local benchmark, we asked whether neighboring-age infant dictionaries can be used interchangeably to represent low-dimensional age-specific scalp patterns. Second, we asked whether a fixed adult-derived cortical basis, mapped onto each infant geometry, could recover infant same-index harmonic coordinates as effectively as the correct age-specific infant basis.

At the neighboring-age level, the answer was already negative. In the main low-dimensional setting of 5 modes per hemisphere, within-age representation was essentially perfect in the noise-free case, whereas representation using a neighboring-age dictionary was systematically degraded. As summarized in Table 1, the symmetrized mismatch loss 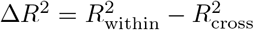 was nonzero for every neighboring-age comparison. The largest losses occurred in several early-to mid-infancy transitions, including 3mo–4.5mo (Δ*R*^2^ ≈ 0.046), 4.5mo–6mo (Δ*R*^2^ ≈ 0.035), and 2mo–3mo (Δ*R*^2^ ≈ 0.028), with mean principal angles between neighboring-age sensor subspaces ranging from roughly 8^◦^ to 21^◦^. Thus, even neighboring-age infant dictionaries are not fully interchangeable for low-dimensional sensor-space representation.

**Table 1:**
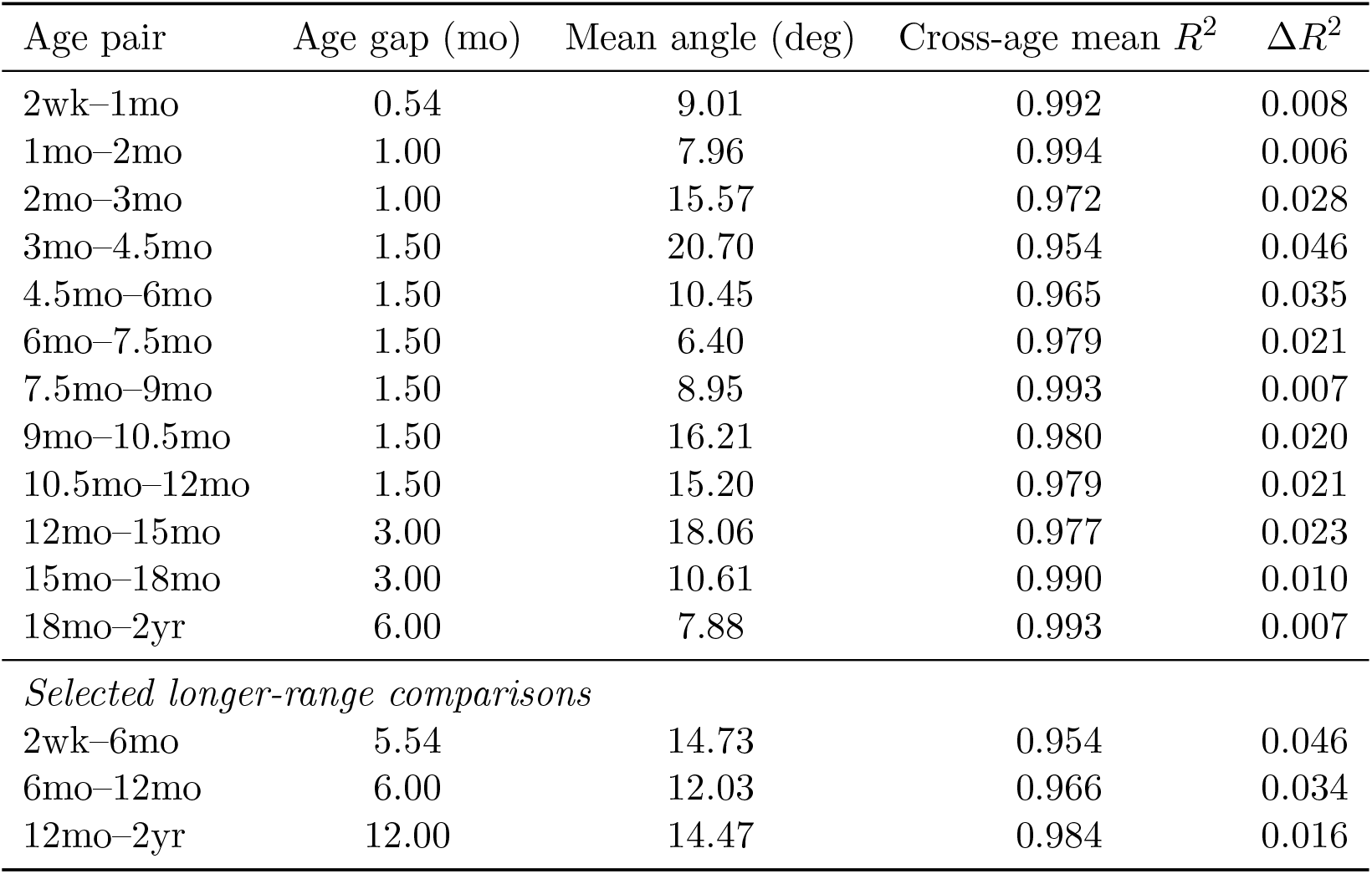
Symmetrized sensor-space mismatch for the main low-dimensional analysis (5 modes per hemisphere). Directional comparisons were symmetrized by averaging *a* → *b* and *b* → *a*, with 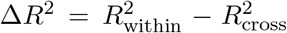. The top block reports neighboring-age comparisons; the bottom block shows selected longer-range comparisons to illustrate that mismatch is not a simple monotone function of age gap.

| Age pair | Age gap (mo) | Mean angle (deg) | Cross-age mean $R^2$ | $\Delta R^2$ |
| --- | --- | --- | --- | --- |
| 2wk–1mo | 0.54 | 9.01 | 0.992 | 0.008 |
| 1mo–2mo | 1.00 | 7.96 | 0.994 | 0.006 |
| 2mo–3mo | 1.00 | 15.57 | 0.972 | 0.028 |
| 3mo–4.5mo | 1.50 | 20.70 | 0.954 | 0.046 |
| 4.5mo–6mo | 1.50 | 10.45 | 0.965 | 0.035 |
| 6mo–7.5mo | 1.50 | 6.40 | 0.979 | 0.021 |
| 7.5mo–9mo | 1.50 | 8.95 | 0.993 | 0.007 |
| 9mo–10.5mo | 1.50 | 16.21 | 0.980 | 0.020 |
| 10.5mo–12mo | 1.50 | 15.20 | 0.979 | 0.021 |
| 12mo–15mo | 3.00 | 18.06 | 0.977 | 0.023 |
| 15mo–18mo | 3.00 | 10.61 | 0.990 | 0.010 |
| 18mo–2yr | 6.00 | 7.88 | 0.993 | 0.007 |
| *Selected longer-range comparisons* |  |  |  |  |
| 2wk–6mo | 5.54 | 14.73 | 0.954 | 0.046 |
| 6mo–12mo | 6.00 | 12.03 | 0.966 | 0.034 |
| 12mo–2yr | 12.00 | 14.47 | 0.984 | 0.016 |

This mismatch becomes more consequential when the analysis basis is fixed at an adult-derived coordinate system (derived from the `fsaverage` cortical surface template; Dale et al., 1999; Fischl et al., 1999; Gramfort et al., 2014). Figure 4 shows that recoverability under the age-specific infant basis (the solid lines) was consistently high and improved with sensor SNR, whereas recoverability under the adult-derived basis (the dotted lines) remained systematically lower across all ages. Thus, a fixed adult-derived basis does not provide an equally effective developmental coordinate system for recovering infant same-index harmonic coordinates.

**Figure 4:**
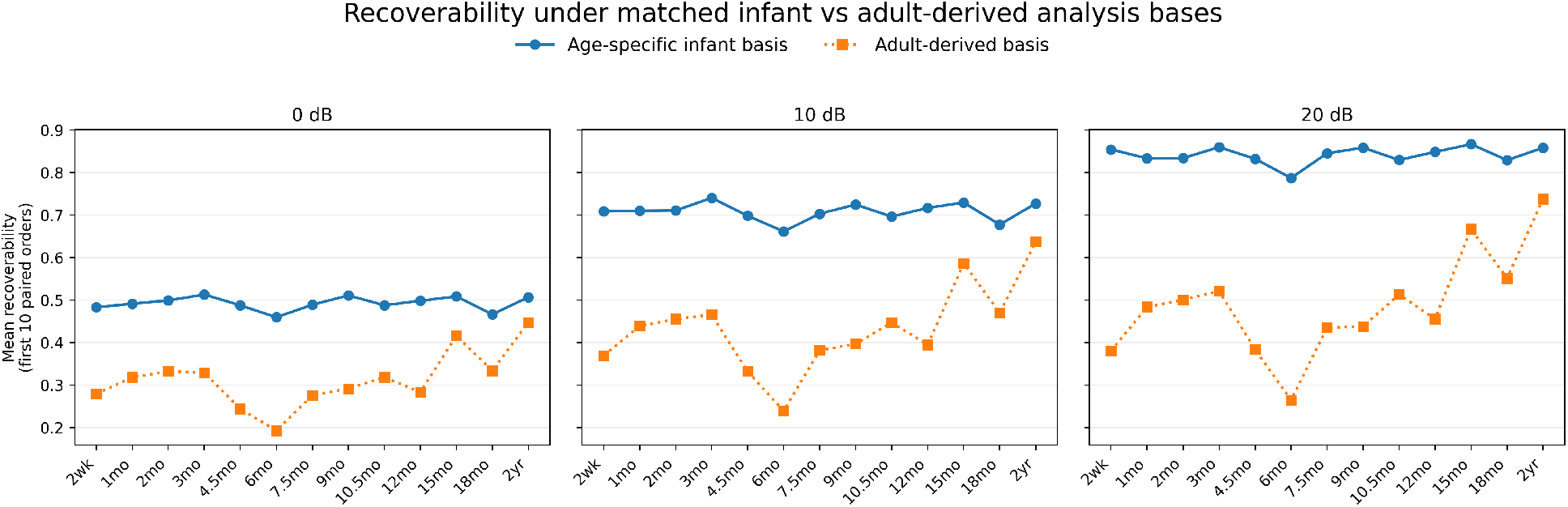
Recoverability under the age-specific infant and adult-derived analysis bases. For each infant age, curves show the mean recoverability of the first 10 paired harmonic orders under two analysis bases: the correct age-specific infant basis (solid blue) and an adult-derived basis mapped to the corresponding infant geometry (dotted orange). Panels correspond to sensor-level SNR values of 0, 10, and 20 dB. Recoverability under the age-specific infant basis is consistently high and improves with SNR, whereas recoverability under the adult-derived basis remains substantially lower across all ages. The gap is especially pronounced in early-to mid-infancy, indicating that a fixed adult-derived cortical basis is systematically suboptimal for recovering infant same-index harmonic coordinates.

At 20 dB, recoverability under the age-specific infant basis was high and relatively stable across age, with mean recoverability rate of the first 10 paired harmonic mode orders ranging from 0.787 to 0.867 (Table 2). In contrast, recoverability under the adult-derived basis varied much more strongly across age, ranging from 0.264 to 0.738. The resulting correct-minus-adult-derived drop ranged from 0.121 to 0.523, with the largest drops at 6mo (0.523), 2wk (0.474), and 4.5mo (0.449), and the smallest drop at 2yr (0.121). Thus, the penalty from using a fixed adult-derived basis tends to be greatest in early-to mid-infancy and becomes smaller, although still nonzero, at later ages.

**Table 2:** Age-by-age recoverability under the age-specific infant and adult-derived analysis bases at 20 dB. Entries report the mean recoverability rate of the first 10 paired harmonic orders. The drop is defined as recoverability under the age-specific infant basis minus recoverability under the adult-derived basis.

| Age | Age-specific infant basis | Adult-derived basis | Drop |
| --- | --- | --- | --- |
| 2wk | 0.855 | 0.380 | 0.474 |
| 1mo | 0.834 | 0.484 | 0.350 |
| 2mo | 0.834 | 0.501 | 0.334 |
| 3mo | 0.860 | 0.521 | 0.339 |
| 4.5mo | 0.833 | 0.384 | 0.449 |
| 6mo | 0.787 | 0.264 | 0.523 |
| 7.5mo | 0.846 | 0.435 | 0.411 |
| 9mo | 0.859 | 0.438 | 0.421 |
| 10.5mo | 0.830 | 0.513 | 0.317 |
| 12mo | 0.849 | 0.456 | 0.393 |
| 15mo | 0.867 | 0.667 | 0.200 |
| 18mo | 0.830 | 0.551 | 0.278 |
| 2yr | 0.859 | 0.738 | 0.121 |

Figure 5 shows that this penalty is also mode-dependent. The recoverability drop is not uniform across the first 10 paired harmonic mode orders. In many ages, the largest drops occur not at the earliest retained modes, but at several intermediate-to-higher retained orders, often around orders 7–10. The age pattern is likewise nonuniform: some ages show broader losses across many retained orders, whereas others show more localized deficits concentrated in a smaller subset of modes. Developmental basis mismatch is therefore not simply a global degradation, but a structured distortion that depends on both age and harmonic mode order.

**Figure 5:**
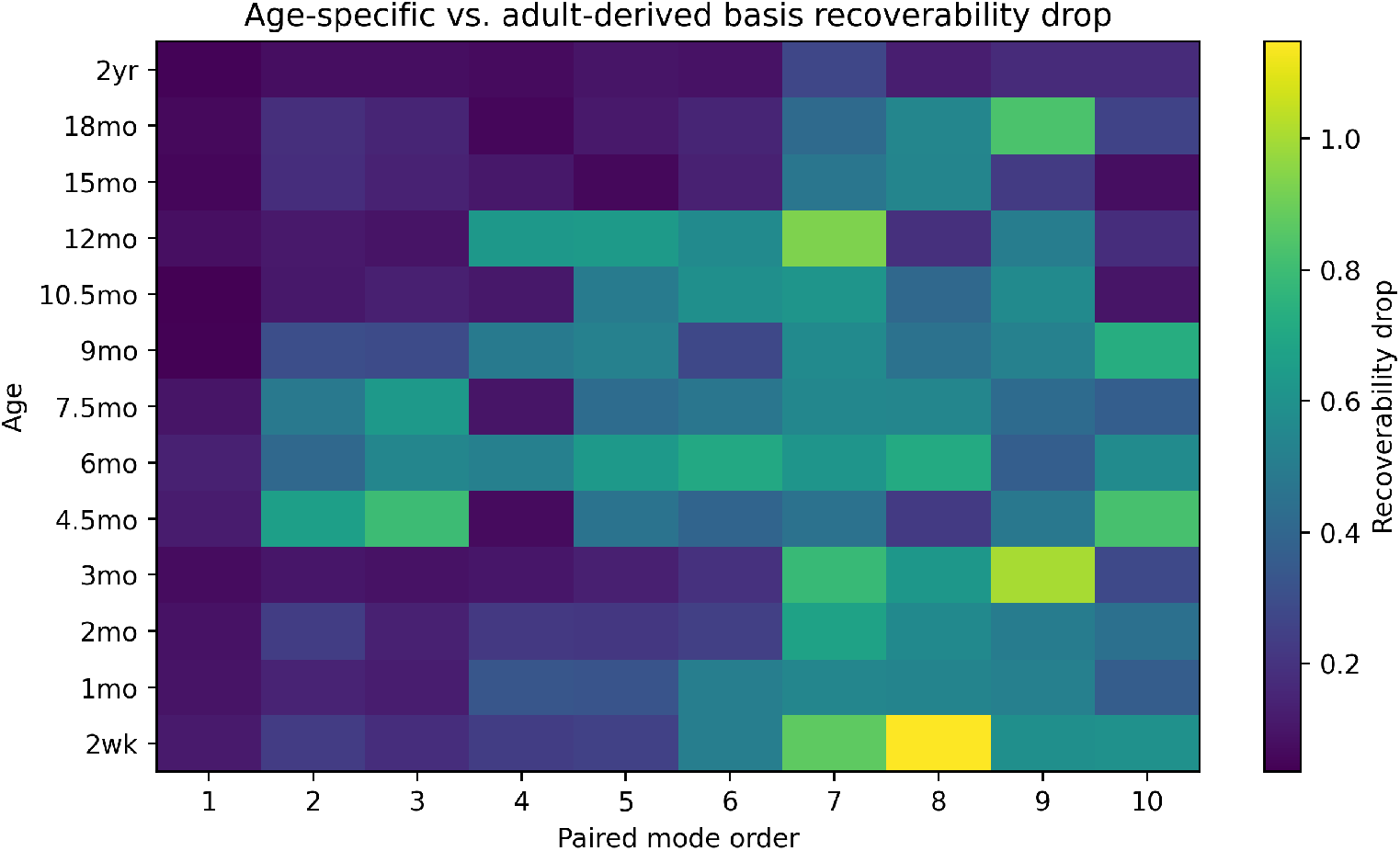
Mode-and age-dependent recoverability penalty from using an adult-derived basis. Heatmap of the correct-minus-adult-derived recoverability drop at 20 dB for each infant age and paired harmonic order. Larger values indicate a greater penalty from replacing the correct age-specific infant basis with the adult-derived basis. The penalty is not uniform across mode order: in many ages it is larger for several intermediate-to-higher retained paired orders than for the earliest retained orders. The age dependence is also nonuniform, with stronger penalties in several early-to mid-infancy templates and smaller penalties at later ages.

These analyses show that developmental basis mismatch has practical consequences in sensor space at both local and broader scales. Neighboring-age infant dictionaries are already not fully interchangeable in low-dimensional sensor space, and a fixed adult-derived basis is systematically suboptimal for recovering infant same-index harmonic coordinates. This pattern supports the view that the appropriate cortical harmonic coordinate system changes across development, motivating age-parameterized or explicitly tracked cortical harmonics rather than a single fixed adult-derived coordinate system.

### 4.4 Neighboring-age cortical harmonic mismatch in basis and coefficient space

We next examined neighboring-age cortical harmonic mode mismatch at two levels: whether neighboring-age cortical LB bases preserve same-index mode correspondence, and whether the coefficients associated with the same cortical pattern remain stable when that pattern is re-expressed in a neighboring-age basis. These analyses show that independently computed neighboring-age cortical harmonic coordinates are only locally comparable by mode index, and that this local basis mismatch is sufficient to destabilize raw coefficient representations.

#### Basis-level neighboring-age mismatch

Figure 6 provides an illustrative example for the neighboring-age pair 1mo and 2mo. In the top row, the comparison is evaluated on the 1mo mesh after mapping the 2mo basis onto 1mo; in the bottom row, it is evaluated on the 2mo mesh after mapping the 1mo basis onto 2mo. The left column shows the left hemisphere and the right column the right hemisphere. In each panel, the dashed diagonal represents ideal same-index correspondence, whereas the cyan trajectory traces the best-matching mapped mode for each evaluation-age mode. The dominant pattern is not complete breakdown, but local off-diagonal displacement: nearby modes often remain related, yet exact same-index correspondence is imperfect.

**Figure 6:**
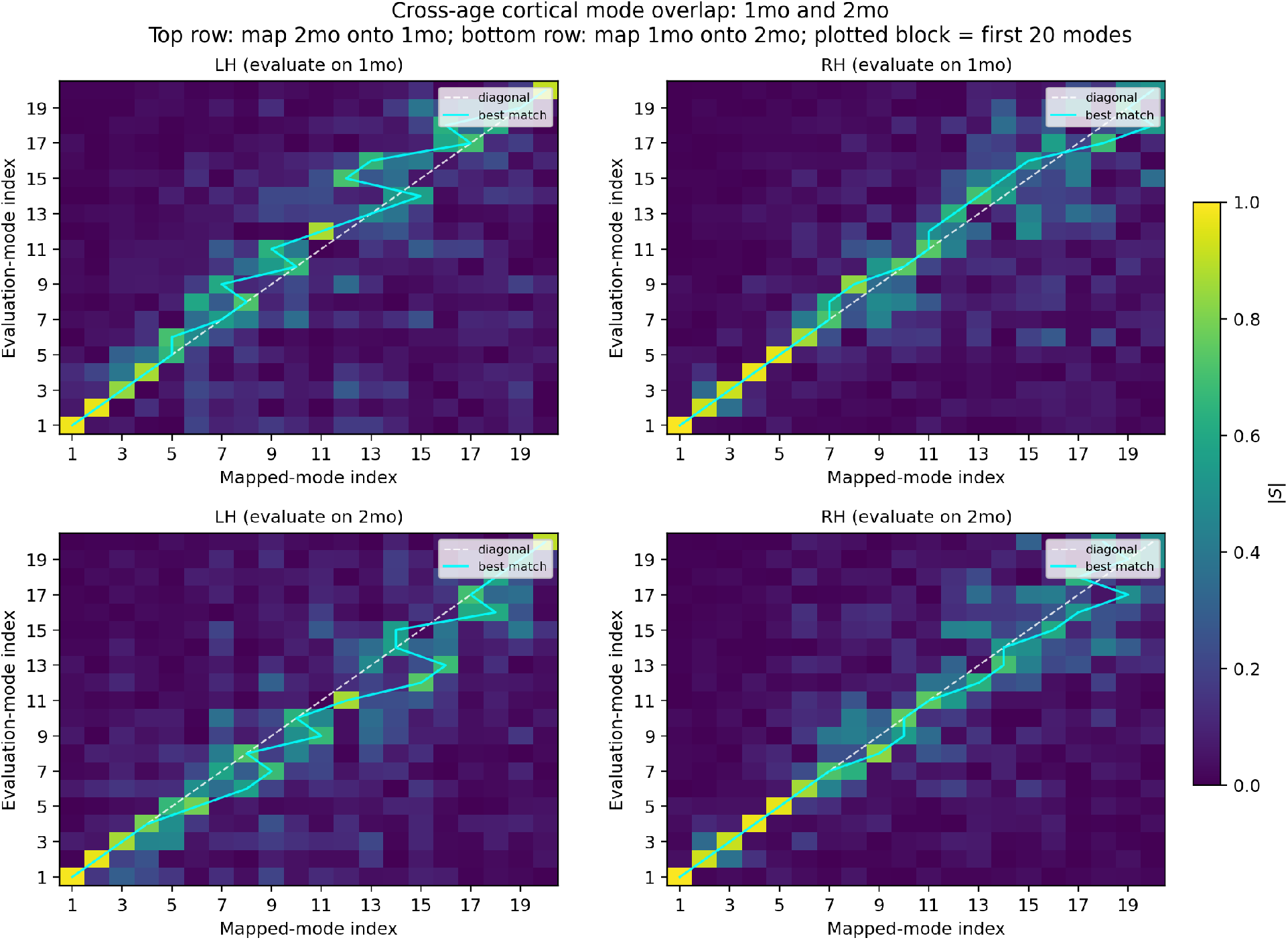
Directional cortical correspondence matrices for the neighboring-age pair 1mo and 2mo. The top row evaluates correspondence on the 1mo mesh after mapping the 2mo basis onto 1mo, and the bottom row evaluates correspondence on the 2mo mesh after mapping the 1mo basis onto 2mo. The left column shows the left hemisphere and the right column the right hemisphere. Rows correspond to evaluation-age mode index and columns to mapped-mode index. The dashed diagonal indicates ideal same-index correspondence, while the cyan curve indicates the best-matching mapped mode for each evaluation-age mode. The full set of neighboring-age directional cortical correspondence matrices is available in the generated output.

Figure 7 summarizes this behavior quantitatively across age. After averaging the two directional comparisons for each neighboring-age pair and then averaging across hemispheres, exact same-index match rates ranged from approximately 0.18 to 0.73, whereas near-neighbor match rates were substantially higher: near-*±*1 rates ranged from approximately 0.53 to 1.00, and near-*±*2 rates from approximately 0.68 to 1.00. Mean absolute best-match shifts ranged from approximately 0.27 to 1.92 mode orders, and adjacent reorderings were consistently nonzero. These results indicate that neighboring-age mismatch is often local rather than catastrophic: low-dimensional harmonic mode organization remains broadly related across age, but individual same-index modes are not reliably preserved, as reflected in the low exact match rates.

**Figure 7:**
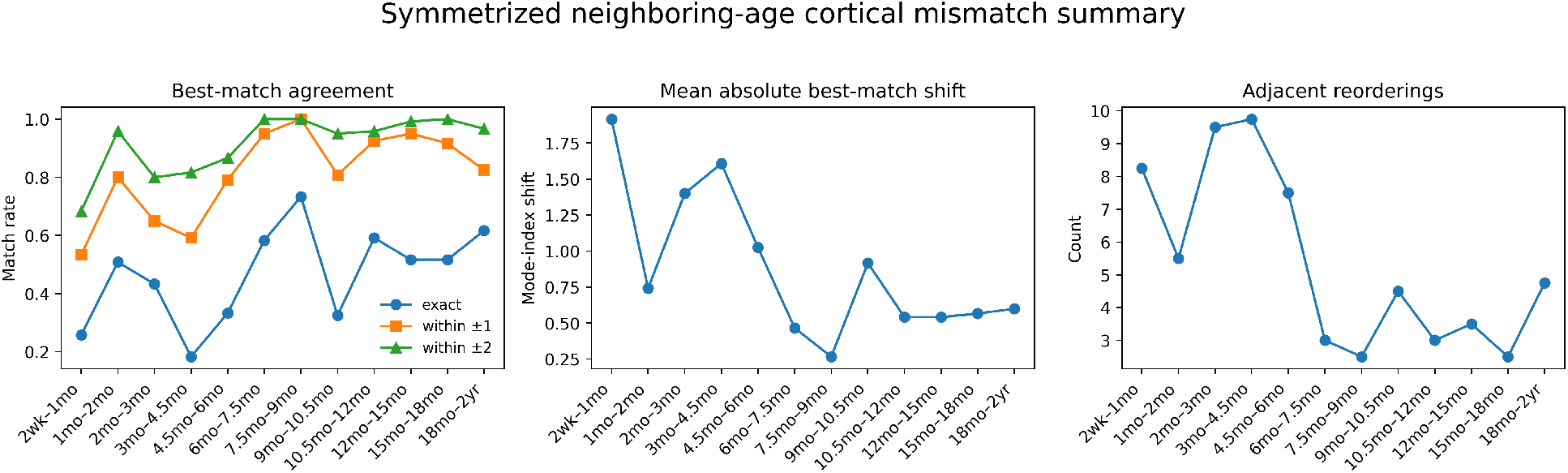
Neighboring-age cortical mismatch summary. For each neighboring-age pair, cortical mismatch metrics were computed in both directions, then averaged across directions and across hemispheres for display. The left panel reports exact and near-neighbor best-match agreement rates, where exact agreement denotes preservation of the same mode index and near-neighbor agreement denotes best matches within *±*1 or *±*2 mode orders. The middle panel shows the mean absolute shift of the best-matching mode index, and the right panel shows the number of adjacent reorderings in the best-match sequence. Across infancy, exact same-index correspondence is often modest, whereas near-neighbor agreement is substantially higher, indicating that developmental mismatch is frequently local rather than catastrophic. At the same time, nonzero mode-index shifts and adjacent reorderings show that independently computed age-specific LB modes are not directly comparable by nominal index alone.

The strongest neighboring-age mismatch occurred in several early-to mid-infancy transitions. In particular, 2wk–1mo and 3mo–4.5mo showed the largest mean shifts (≈ 1.92 and ≈ 1.61, respectively) (middle panel), and 3mo–4.5mo also had the lowest exact same-index agreement (≈ 0.18) (first panel). By contrast, several later intervals showed more stable local correspondence. For example, 7.5mo–9mo showed the highest exact same-index agreement (≈ 0.73) and the smallest mean shift (≈ 0.27), while 6mo–7.5mo and 10.5mo–12mo also showed relatively small shifts and high near-neighbor agreement. Thus, independently computed same-index cortical LB modes are not directly comparable across development without explicit age-parameterization or tracking, even though neighboring-age harmonic subspaces often remain locally related.

#### Coefficient-level neighboring-age mismatch

Figure 8 shows the downstream consequence of this basis-level non-equivalence. Rather than asking whether neighboring-age bases are aligned, we now ask how much the coefficients associated with the same cortical pattern change when expressed in neighboring-age bases. For the *single-mode test patterns* (top panels), the fraction of squared target coefficient mass remaining at the same nominal mode index after re-expression in the neighboring-age basis was often small-to-modest, ranging from 0.172 to 0.560. However, substantially more squared coefficient mass remained within nearby modes: the fraction retained within *±*1 neighboring indices ranged from 0.434 to 0.848, and the fraction retained within *±*2 ranged from 0.557 to 0.935. Thus, neighboring-age non-equivalence rarely preserves the same nominal mode exactly, but much of the coefficient mass remains concentrated in a nearby neighborhood of modes. The strongest mismatches again appeared in several early-to mid-infancy transitions, especially 2wk–1mo, 2mo–3mo, 3mo–4.5mo, and 9mo–10.5mo, whereas 6mo–7.5mo and 7.5mo–9mo were among the most stable neighboring intervals.

**Figure 8:**
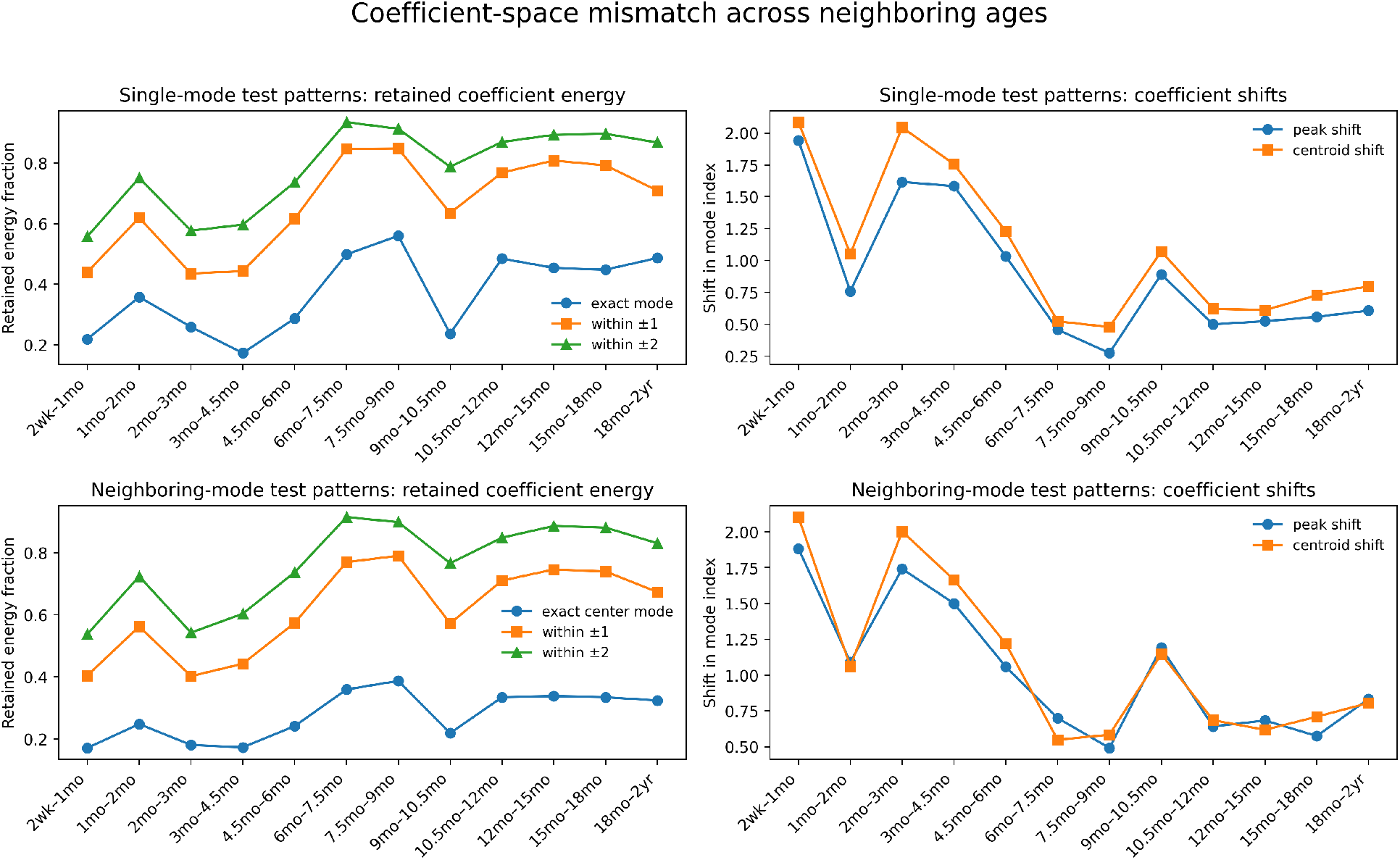
Coefficient-space mismatch across neighboring ages. Directional comparisons were averaged across the two mapping directions and then across hemispheres. The top row shows results for *single-mode test patterns*, in which a cortical field is generated from one nominal harmonic mode and then re-expressed in the basis of a neighboring age. The bottom row shows results for *neighboring-mode test patterns*, in which the source-age coefficient vector is concentrated in a small neighborhood of nearby mode indices rather than at a single mode. For both pattern families, the left column reports the fraction of squared target coefficient mass remaining at the same nominal mode index (*exact*) and within *±*1 and *±*2 neighboring mode indices. The right column reports coefficient shifts: the mean absolute displacement of the peak target coefficient index (*peak shift*) and of the energy-weighted center of mass of the target coefficient distribution (*centroid shift*). Across infancy, neighboring-age non-equivalence generally causes squared coefficient mass to leak into nearby modes rather than remaining at the same nominal index, and this local leakage is sufficient to make raw age-specific coefficient representations unstable for longitudinal use.

The *neighboring-mode test patterns* (bottom panels) showed the same qualitative behavior, but with slightly more diffuse concentration, as expected for patterns that already occupy a small local band of mode indices rather than a single exact mode. The fraction of squared target coefficient mass remaining at the same nominal center mode ranged from 0.171 to 0.387, whereas the fraction retained within *±*1 neighboring indices ranged from 0.403 to 0.791, and the fraction retained within *±*2 ranged from 0.538 to 0.915. Thus, even when the source-age pattern is already distributed over a small local neighborhood, re-expression in a neighboring-age basis still produces substantial local leakage and mode-index drift.

Taken together, Figures 6, 7, and 8 show a coherent pattern. Independently computed neighboring-age cortical harmonic bases retain substantial local structure, but not strict same-index correspondence. This local basis-level mismatch propagates directly to the coefficient level as leakage of squared coefficient mass into nearby modes and instability of raw coordinate representations. Neighboring-age cortical harmonic coordinates are therefore not globally unrelated, but they are not stable enough by nominal mode index to support straightforward longitudinal use without explicit age-parameterization or tracking.

### 4.5 Modes in more crowded local eigenvalue neighborhoods are harder to match across age

We next asked whether cortical cross-age instability is concentrated in parts of the ordered basis where a mode’s eigenvalue lies unusually close to its neighbors. Figure 9 summarizes this relationship using a crowding score derived from the relative gap between each mode’s eigenvalue and the closest neighboring eigenvalue in the ordered spectrum. The left panel shows substantial variation in this crowding score across both age and mode order, indicating that some portions of the basis are much more tightly packed than others.

**Figure 9:**
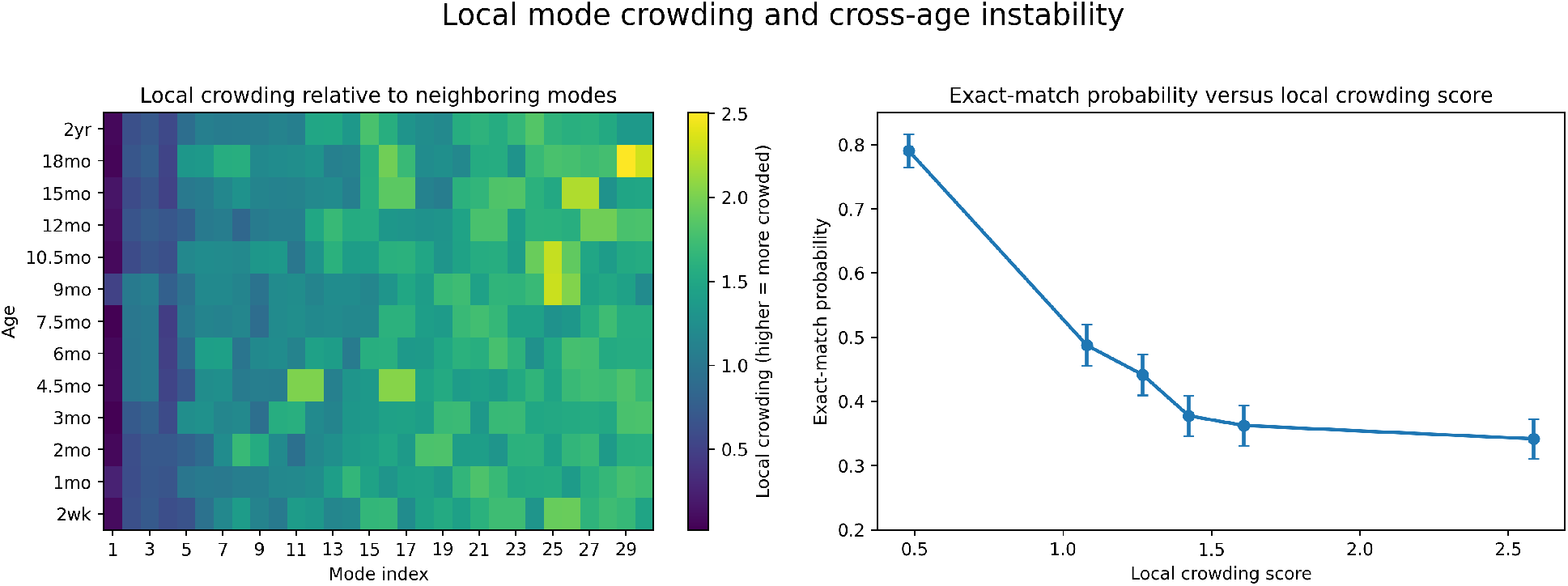
Local eigenvalue crowding and cross-age instability. Left: age-by-mode-order heatmap of the crowding score, which is larger when a mode’s eigenvalue lies closer to its nearest neighboring eigenvalue in the ordered basis. Right: exact same-index match probability as a function of the crowding score, obtained by pooling modes across ages and grouping them into bins of similar crowding. Modes in more crowded neighborhoods are less likely to preserve exact same-index correspondence across age, indicating that cross-age non-equivalence occurs preferentially where nearby modes are packed more tightly together.

The right panel of Figure 9 shows how exact same-index matching depends on this crowding score. For each neighboring-age comparison, hemisphere, and evaluation-age mode, we linked that mode’s crowding score to whether its best match remained at the same nominal index, and then grouped these mode-level observations into bins of similar crowding. The exact same-index matching probability was clearly less likely in more crowded neighborhoods. In other words, modes whose eigenvalues lie especially close to neighboring eigenvalues are harder to match exactly across age. A natural interpretation is that when local eigenvalue spacing is small, modest developmental changes can more easily induce local reordering or rotation among nearby modes. Failures of strict same-index matching are therefore not arbitrary: they occur preferentially where the local mode sequence is most crowded. This supports the idea that cross-age comparison should be neighborhood-aware or subspace-aware rather than strictly indexwise.

### 4.6 Both cortical-basis and forward-operator changes contribute to neighboring-age sensor-space mismatch

We next examined whether neighboring-age sensor-space mismatch is more strongly associated with changes in the cortical eigenmode basis or with changes on the forward-operator side. To do so, we compared the *full neighboring-age mismatch*—that is, the mismatch between the two native age-specific sensor dictionaries for a given neighboring-age pair—with two partial substitution analyses: one in which only the cortical basis was replaced while the forward operator was held fixed, and one in which only the forward-operator side was replaced while the cortical basis was held fixed.

As shown in Figure 10, both cortical-basis and forward-operator-side changes contributed to neighboring-age sensor-space mismatch. Note that these are one-at-a-time substitution comparisons rather than additive decompositions of total mismatch. Across neighboring-age pairs, the forward-operator-side substitution generally produced mismatch values closer to the full neighboring-age mismatch than did the cortical-basis substitution. Consistent with this, the forward-operator-side substitution exceeded the cortical-basis substitution in 8*/*12 neighboring pairs for Δ*R*^2^ and in 7*/*12 pairs for projector distance (Table 3). Median Δ*R*^2^ values were 0.0200 for the full neighboring-age mismatch, 0.0065 for the cortical-basis substitution, and 0.0091 for the forward-operator-side substitution; the corresponding median projector distances were 1.550, 1.036, and 1.112.

**Table 3:** Summary of partial substitution analyses for neighboring-age sensor-space mismatch. Values are medians across neighboring-age pairs. The cortical-basis substitution uses the neighboring-age cortical basis while holding the forward operator fixed. The forward-operator-side substitution uses the neighboring-age forward-operator side while holding the cortical basis fixed.

| Metric | Full neighboring-age mismatch | Cortical-basis substitution | Forward-operator-side substitution |
| --- | --- | --- | --- |
| $\Delta R^2$ (drop due to mismatch) | 0.0200 | 0.0065 | 0.0091 |
| Projector distance | 1.550 | 1.036 | 1.112 |

**Figure 10:**
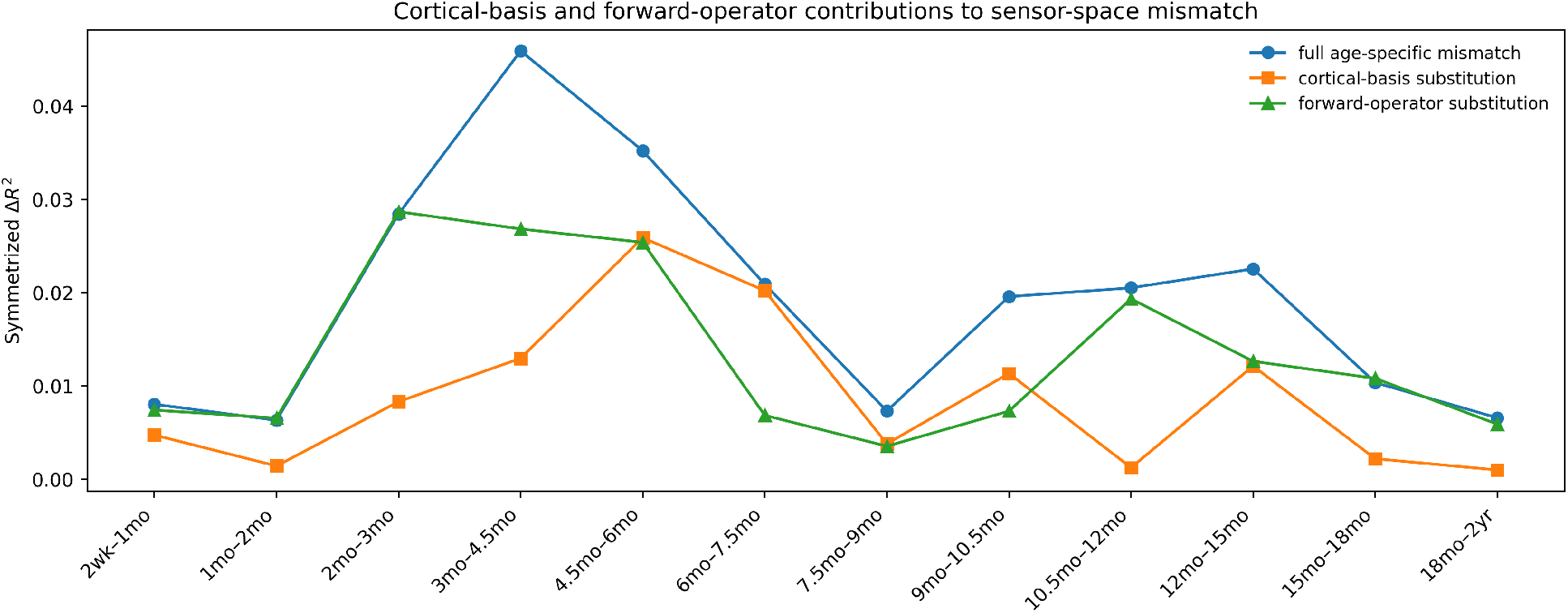
Cortical-basis and forward-operator-side contributions to neighboring-age sensor-space mismatch. For each neighboring-age pair, the blue curve shows the full neighboring-age mismatch between the two native age-specific sensor dictionaries. The orange curve shows the cortical-basis substitution, in which the cortical basis is replaced while the forward operator is held fixed. The green curve shows the forward-operator-side substitution, in which the forward-operator side is replaced while the cortical basis is held fixed. Across many neighboring-age pairs, the forward-operator-side substitution is comparable to or larger than the cortical-basis substitution, although cortical-basis change remains substantial in several mid-infancy intervals.

At the same time, cortical-basis mismatch remained substantial in several intervals, especially during mid-infancy, and in some comparisons was comparable to or larger than the forward-operator-side contribution. In particular, the cortical-basis substitution was similar to or larger than the forward-operator-side substitution in intervals such as 4.5mo–6mo, 6mo–7.5mo, and 9mo–10.5mo. The largest full neighboring-age mismatch occurred in early-to mid-infancy, most notably around 2–6 months. Overall, these results show that neighboring-age low-dimensional sensor mismatch is not explained by cortical-basis change alone: changes on the forward-operator side are often at least as important, but cortical-basis change remains substantial in several mid-infancy transitions.

### 4.7 Sequential Procrustes tracking improves same-index neighboring-age consistency

Sequential Procrustes tracking substantially improved same-index neighboring-age consistency of the cortical harmonic basis. Figure 11 shows that exact-match rate increased for all neighboring-age pairs after tracking, and mean absolute best-match shift decreased in most intervals. After averaging over both directional evaluations and the two hemispheres, and then summarizing across the 12 neighboring-age pairs, the median exact-match rate for the first 30 modes increased from 0.513 before tracking to 0.825 after tracking, while the corresponding median mean absolute best-match shift decreased from 0.671 to 0.529 mode orders. Thus, the tracking procedure consistently improved preservation of same-index mode identity across age.

**Figure 11:**
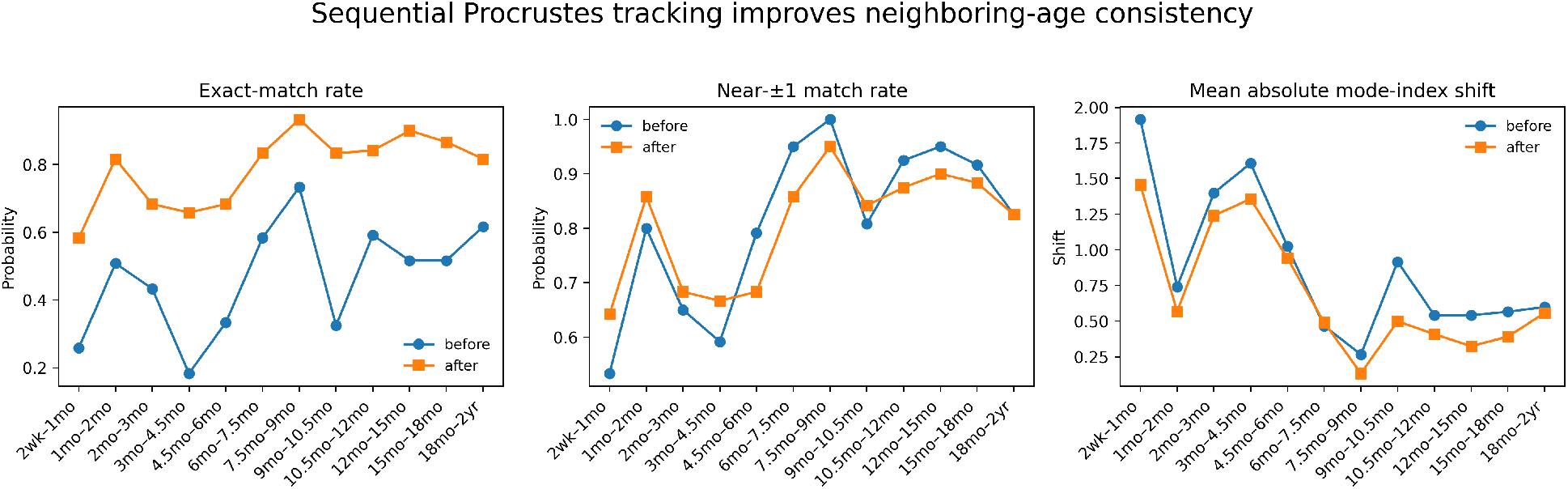
Sequential Procrustes tracking improves neighboring-age harmonic consistency. For each neighboring-age pair, curves compare overlap-based consistency summaries before and after Sequential Procrustes tracking of the cortical harmonic basis. Left: exact same-index match rate. Middle: near-*±*1 match rate. Right: mean absolute best-match shift in mode index. Sequential Procrustes tracking substantially improves exact same-index correspondence across neighboring ages and reduces best-match shift in most intervals. Changes in near-neighbor match rate are less uniform because many raw neighboring-age comparisons already exhibited substantial neighborhood-level agreement before tracking; the main effect of the procedure is to sharpen those local correspondences into more exact same-index matching. The resulting basis functions should be interpreted as *Procrustes-tracked harmonic basis functions* rather than as exact tracked eigenvectors, because blockwise rotations generally move the columns away from strict Laplace–Beltrami eigenfunctions while preserving the relevant low-dimensional harmonic subspaces.

The improvement was especially pronounced in developmental intervals that previously exhibited strong local reordering. For example, exact-match rates increased markedly for intervals such as 2wk–1mo, 3mo– 4.5mo, 9mo–10.5mo, 12mo–15mo, and 15mo–18mo. Even in intervals that were already relatively stable before tracking, such as 6mo–7.5mo and 7.5mo–9mo, exact-match rates increased further and approached ceiling values. Overall, tracking increased exact-match rate in 12*/*12 neighboring-age pairs and reduced mean absolute best-match shift in 11*/*12 comparisons.

Near-neighbor match rates changed less uniformly. This is expected because many raw neighboring-age comparisons already exhibited substantial neighborhood-level agreement before tracking. The main effect of Sequential Procrustes tracking is therefore not to increase broad neighborhood-level agreement, but to sharpen those local correspondences into more exact same-index alignment.

Taken together, these results show that a substantial portion of the raw neighboring-age mismatch can be reduced by explicit local subspace alignment. This suggests that much of the instability of independently computed neighboring-age harmonic bases arises from local rotations and reorderings within nearby harmonic neighborhoods rather than from complete loss of shared low-dimensional structure. As a down-stream validation, Sequential Procrustes tracking also increased exact same-index coefficient retention for both single-mode and neighboring-mode test patterns across all neighboring-age pairs (Supplementary Figure S3).

## 5 Discussion

Our results support a clear developmental conclusion: across infancy, developmental change affects both what scalp EEG can express and how cortical harmonic coordinates should be compared across age. On the one hand, age-specific forward models consistently emphasized lower-order cortical harmonics at the scalp, and the relative coarse-to-fine transfer profile was broadly stable across infancy. On the other hand, practical coordinate recovery depended not only on noise and dimensionality, but also on whether the analysis basis matched developmental stage: a fixed adult-derived basis was systematically suboptimal for recovering infant same-index harmonic coordinates. In this sense, the present study extends recent mature-brain eigenmode work by showing that a single fixed cortical basis is not sufficient when cortical geometry changes rapidly across development (Müller et al., 2022; Pang et al., 2023; Park, 2026).

The cross-age analyses sharpen this point. In cortical mode space, independently computed LB bases were not directly comparable by nominal index, because neighboring ages exhibited local mode mixing, reordering, and index drift. In sensor space, this non-equivalence was not merely a mathematical feature of the cortical eigensystem: age-mismatched dictionaries produced measurable loss of low-dimensional representation quality. At the coefficient level, even when squared coefficient mass remained concentrated within a nearby neighborhood of modes, raw coefficients were still unstable across age. Together, these findings argue against naive same-index longitudinal comparison and instead support age-parameterized or explicitly tracked cortical harmonic coordinates for developmental EEG.

The supporting analyses refine that conclusion. The local-spacing analysis showed that exact same-index instability was concentrated in crowded parts of the ordered spectrum, where nearby eigenvalues are more tightly spaced and therefore more susceptible to local rotation or reordering under developmental change. The partial substitution analysis further showed that neighboring-age sensor-space mismatch was not explained by cortical-basis change alone: changes on the forward-operator side were often at least as important, although cortical-basis change remained substantial in several mid-infancy transitions. This distinction is important. Age-parameterized cortical harmonics are needed to stabilize the cortical coordinate system itself, but full sensor-space comparability also depends on age-varying forward-model effects. This interpretation is consistent with recent work showing that scalp spatial structure is shaped jointly by cortical organization and realistic head geometry (Srinivasan et al., 1996, 1998; Nunez and Srinivasan, 2006; Graichen et al., 2025).

Sequential Procrustes tracking provides a useful local alignment remedy. It substantially improves exact same-index neighboring-age consistency and same-index coefficient retention, showing that a large portion of the observed neighboring-age instability is due to local subspace misalignment rather than complete loss of shared low-dimensional structure. At the same time, this tracking procedure is intentionally modest: for example, it does not jointly estimate a smooth age-dependent basis trajectory across all ages, nor does it include subject-specific deviations or random effects around such a trajectory. In that sense, it is best interpreted as evidence that explicit alignment helps, while also motivating the need for a more principled age-parameterized harmonic model.

Because the present study is template-based, its primary value is mechanistic rather than subject-specific. It isolates how developmental changes in cortical geometry and head conduction shape harmonic observability and cross-age coordinate comparability in principle. This template-based perspective is especially relevant in infancy, where subject-specific MRI is often unavailable and template-based source modeling remains an important practical strategy (Richards et al., 2016; O’Reilly et al., 2021). The main current limitations are the use of age-specific templates rather than subject-specific anatomical models, the reliance on a sparse template 10–20 montage in the primary analyses, and the absence of a fully statistical age-parameterized tracking model. These limitations point naturally toward future work: validation with subject-specific anatomy and denser montages (Asayesh et al., 2024; Dickinson et al., 2024), explicit developmental modeling of harmonic subspaces, and practical extensions that connect template-based developmental analysis to MRI-free electromagnetic source-imaging workflows (He et al., 2018; O’Reilly et al., 2021; Jaiswal et al., 2025).

## 6 Conclusion

Across infancy, age-specific forward models preferentially transmit lower-order cortical harmonics to the scalp, but developmental changes in cortical geometry alter how these harmonics should be indexed and compared across age. Independently computed age-specific cortical LB modes are only locally comparable by nominal index, and this local mismatch is sufficient to produce measurable instability in low-dimensional sensor-space representation and coefficient-space coordinates. A fixed adult-derived basis is systematically suboptimal for recovering infant same-index harmonic coordinates, whereas explicit local tracking substantially improves same-index consistency across neighboring ages. Together, these findings motivate age-parameterized or explicitly tracked cortical harmonic coordinates as a principled foundation for longitudinal developmental EEG and, more broadly, for lifespan neuroimaging analyses that seek geometry-aware spatial coordinates across changing brains.

## Data availability

The age-specific infant anatomical templates used in this study are publicly available (through MNE-Python).

## Supplementary Materials

### S1 Details of age-specific cortical Laplace–Beltrami eigenmode computation

For each template age, cortical Laplace–Beltrami eigenmodes were computed separately on the left-and right-hemisphere template white surfaces. Let the cortical mesh at age *a* be represented by vertices and triangular faces. We used the standard finite-element discretization of the Laplace–Beltrami operator, yielding the generalized eigenvalue problem

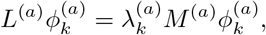

where *L*^(*a*)^ is the cotangent Laplacian and *M* ^(*a*)^ is the area-weighting mass matrix (Meyer et al., 2003; Reuter et al., 2006). The eigenvalues were ordered increasingly, so that smaller 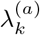 correspond to spatially smoother cortical modes and larger 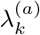 correspond to spatially finer ones.

The eigenvectors were normalized with respect to the area-weighted inner product induced by *M* ^(*a*)^, so that

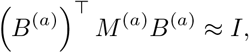

Where 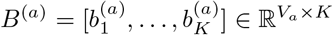. As usual for Laplace-type eigenproblems, each eigenvector is defined only up to an arbitrary global sign, and modes in closely spaced local neighborhoods of the spectrum can be unstable to local rotations or reorderings across age. These properties are central to the cross-age comparability questions studied in the main text.

For forward-model analyses, the full-mesh eigenvectors were then restricted to the age-specific EEG source-space vertices used by the leadfield matrix. This restriction is necessary because the forward model is defined on the source-space discretization rather than on the full cortical mesh. After restriction, each source-space basis column was re-normalized to unit Euclidean norm before constructing the forward-projected dictionary. This ensures that age-to-age differences in forward-projected column norms reflect the age-specific forward model and cortical geometry rather than trivial differences in source-space vertex count or source pruning.

### S2 Details of spherical-registration mapping for cross-age cortical comparisons

For cortical correspondence and coefficient-space transfer analyses, we map cortical quantities between ages using the template spherical-registration surfaces provided with the infant templates (Dale et al., 1999; Fischl et al., 1999). Let

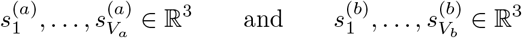

denote the spherical-registration coordinates of the age-*a* (with *V*_*a*_ number of vertices) and age-*b* (with *V*_*b*_ number of vertices) meshes. After normalizing these spherical coordinates to unit length, for each target vertex *v* on age *b* we identify the nearest source vertex on age *a*,

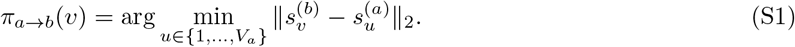

We then transfer values by nearest-neighbor interpolation on the registered sphere. This gives a topology-aware mapping operator for the present template-based comparisons and ensures that all cross-age cortical correspondence and coefficient-transfer analyses are defined on the target mesh.

### S3 Details of sensor-space representation metrics

For the sensor-space mismatch analysis in Section 3.3, we compare within-age and cross-age representation of reference-age patterns under low-dimensional sensor subspaces. Let *Q*_*a*_ and *Q*_*b*_ denote orthonormal bases for span(*D*^(*a*)^) and span(*D*^(*b*)^), respectively. We evaluate representation quality using mean-centered sensor-space *R*^2^, computed after subtracting the across-channel mean from both the observed pattern and its projection. This avoids trivial inflation of fit quality due to shared constant offsets across channels.

We additionally compute principal angles (Knyazev and Argentati, 2002) between the two sensor sub-spaces from the singular values of 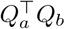, and summarize subspace distance by the Frobenius norm of the difference between the corresponding projectors,

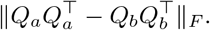

When the analyzed subspace approaches the full sensor dimension, these subspace-comparison metrics can become artificially small because the sensor space is nearly saturated. This is why the main analysis focuses on 5 modes per hemisphere.

### S4 Detailed formulation of coefficient-space mismatch across age

The cross-age correspondence matrix quantifies basis alignment, but our eventual goal is longitudinal modeling in terms of harmonic *coefficients*. Accordingly, we define a coefficient-space mismatch analysis that asks how much the coordinates of the same underlying cortical pattern change when expressed in age-specific harmonic bases at different ages.

#### Age-specific harmonic bases

Let 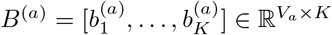 denote the cortical harmonic basis at age *a*, where *V*_*a*_ is the number of vertices on the age-*a* cortical mesh and *K* is the number of retained modes. In the current untracked analysis, *B*^(*a*)^ is the age-specific LB basis on the full cortical mesh. In the Sequential Procrustes tracked analysis, the same formulation is reused by replacing *B*^(*a*)^ with a tracked basis 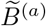.

#### Within-age coefficients

For a cortical field at age *a*,

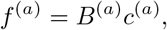

the within-age coefficient vector is obtained by projection:

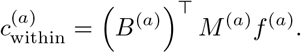

The matrix *M* ^(*a*)^ is needed because the natural inner product between two cortical fields is the surface *L*^2^ inner product,

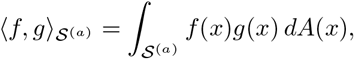

which is approximated on the mesh by

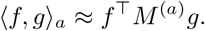

Thus, projection onto the harmonic basis, cortical correspondence calculations, and coefficient transfer all use the area-weighted inner product rather than an unweighted Euclidean dot product.

#### Mapping a field from age *a* **to age** *b*

To compare the same field across age, we first transport it from the cortical mesh at age *a* to the cortical mesh at age *b* using the template spherical registration. Each age-specific cortical template includes a spherical-registration surface (`sphere.reg`) that maps the cortical mesh to a common spherical coordinate system while preserving topological correspondence as well as possible.

For each target vertex *v* on age *b*, we identify the nearest source vertex on age *a, π*_*a*→*b*_(*v*) in (S1). We then define the transported field by nearest-neighbor interpolation on the registered sphere:

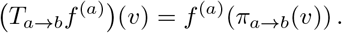

#### Cross-age coefficients

Once the field *f* ^(*a*)^ has been transported to the target mesh, its coefficients in the age-*b* basis are

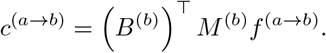

If the two age-specific bases were perfectly comparable, then *c*^(*a*→*b*)^ would match *c*^(*a*)^ up to sign, permutation, or explicit alignment. Departures from this ideal represent coefficient-space mismatch.

#### Coefficient transfer operator

The entire mapping from source-age coefficients to target-age coefficients can be written as

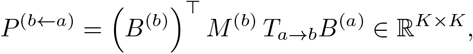

so that

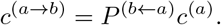

If *P* ^(*b*←*a*)^ were close to the identity matrix, then same-index coefficients would be stable across age. Offdiagonal structure in *P* ^(*b*←*a*)^ indicates coefficient mixing.

#### Families of test patterns

We use two families of coefficient vectors to probe different aspects of coordinate stability.

##### 1 Single-mode test patterns

We set

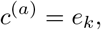

so that the resulting field is exactly the *k*-th harmonic mode at age *a*. This isolates how an individual nominal mode spreads under age mismatch.

##### 2. Neighboring-mode test patterns

We define a Gaussian-weighted coefficient “packet” centered at mode *k*,

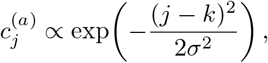

with *σ* = 1 mode-index unit in the reported analyses, followed by normalization. These patterns test whether a cortical pattern concentrated in a small neighborhood of mode indices remains concentrated in a nearby neighborhood after age mismatch.

#### Summary metrics

For the single-mode and neighboring-mode test patterns, we summarize coefficient mismatch by: (i) the fraction of coefficient energy retained at the nominal source mode, (ii) the fraction retained within *±*1 and *±*2 neighboring mode indices, (iii) the shift in the highest-energy target coefficient index, and (iv) the energy-weighted centroid shift of the coefficient mass.

#### Relation to tracking

This formulation is designed to be reusable for tracked-basis analyses. Once a tracked basis 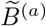 is available, the same code and summary metrics can be reused by substituting 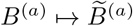, allowing direct comparison between independently computed coordinates and explicitly aligned coordinates.

### S5 Implementation details of the adult-basis mismatch recoverability analysis

The “correctly matched” recoverability analysis uses the infant age-specific analysis dictionary

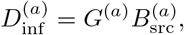

where 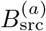 is the infant cortical harmonic basis restricted to the age-*a* EEG source space and *G*^(*a*)^ is the corresponding infant forward operator.

For the adult-basis mismatch analysis, we begin with a standard adult cortical harmonic basis *B*_adult_, computed on the `fsaverage` cortical surface template (Dale et al., 1999; Fischl et al., 1999; Gramfort et al., 2014). This adult basis is mapped to each infant age-*a* cortical mesh using the same spherical-registration mapping described in Supplementary Section S2. After mapping, the adult basis is restricted to the age-*a* EEG source space and each column is re-normalized to unit Euclidean norm. The resulting source-space adult-derived basis is then sign-aligned columnwise to the age-*a* infant basis by nominal mode index, so that each adult-derived column has positive correspondence with the infant column of the same nominal order. This removes arbitrary global sign ambiguity while preserving the substantive developmental basis mismatch. The resulting sign-aligned adult-derived basis is denoted

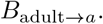

The corresponding adult-derived analysis dictionary is then

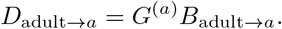

Because this dictionary uses the infant forward operator *G*^(*a*)^, differences between the correctly matched and adult-mismatch analyses reflect mismatch in the cortical analysis basis rather than mismatch in the forward model.

For each infant age *a*, latent infant-basis coefficients *w*^(*a*)^(*t*) are simulated and noisy scalp observations are generated from

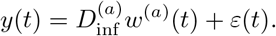

Thus, the latent cortical pattern is always generated in the infant age-specific coordinate system. In the matched analysis, coefficients are recovered using *D*^(*a*)^; in the adult-mismatch analysis, coefficients are recovered using *D*_adult→*a*_.

The recovered coefficients in the adult-mismatch analysis are compared directly with the true infant coefficients *w*^(*a*)^(*t*) by the same nominal mode index, after the sign-alignment step described above. This is intentionally an index-based developmental mismatch analysis designed to ask the following question: if one insists on using a fixed adult-derived cortical coordinate system, how well can one recover the infant harmonic coordinates of the same nominal order? Under this design, any degradation reflects the combined effect of realistic sensor noise and developmental mismatch between the infant generating basis and the adult-derived analysis basis.

The primary summary of interest is therefore the drop in recoverability relative to the correctly matched infant-basis benchmark. A large drop indicates that the adult-derived basis fails to recover infant same-index harmonic coordinates, supporting the need for age-parameterized or explicitly tracked cortical coordinates in developmental EEG.

### S6 Implementation details of Sequential Procrustes tracking

Sequential Procrustes tracking was implemented separately for the left and right hemispheres. Let

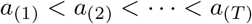

denote the ordered template ages, and let 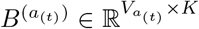 denote the corresponding age-specific cortical LB eigenmode basis. The youngest age served as the initial reference, so that

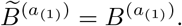

For each subsequent age *a*_(*t*+1)_, we first mapped the native basis 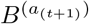 onto the cortical mesh of age *a*_(*t*)_ using the spherical-registration mapping operator described in Supplementary Section S2. This yielded a mapped basis

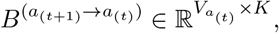

defined on the same mesh as the previously tracked basis 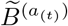.

To reflect the fact that most neighboring-age mismatch appeared local rather than global, we partitioned the first *K* modes into small contiguous groups of nearby indices. In the current implementation, we used groups of three consecutive modes, for example {1, 2, 3}, {4, 5, 6}, and so on. Let ℐ_*g*_ denote one such group, and write

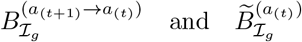

for the mapped current-age basis and the previously tracked basis restricted to that same group of columns.

For each group, we solved the weighted orthogonal Procrustes problem

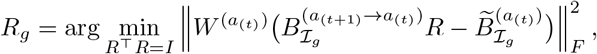

where

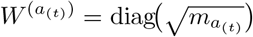

is the diagonal matrix of square-root vertex-area weights on the age-*a*_(*t*)_ mesh. This weighting makes the alignment objective consistent with the area-weighted inner product used throughout the cortical correspondence analysis.

The resulting orthogonal matrix *R*_*g*_ was then applied to the corresponding native group of modes at age *a*_(*t*+1)_:

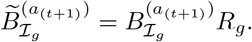

After all groups had been rotated in this way, we applied a columnwise sign adjustment. Specifically, each tracked column at age *a*_(*t*+1)_ was mapped back to the age-*a*_(*t*)_ mesh, and its sign was flipped if necessary so that its area-weighted inner product with the corresponding previously tracked column at age *a*_(*t*)_ was positive. This removes the arbitrary sign ambiguity of the eigenmodes and makes the sequential alignment consistent across ages.

The final output of this procedure is a sequence of Procrustes-tracked bases

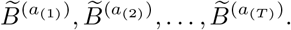

Because the rotations are applied within local groups of nearby modes, the resulting columns generally no longer coincide exactly with the original LB eigenvectors. The procedure should therefore be interpreted as producing a locally aligned cortical eigenmode basis rather than a strict tracked eigensystem.

### S7 Hemisphere-specific cortical mismatch summaries

In the main text, neighboring-age cortical correspondence summaries are averaged across hemispheres to emphasize the overall developmental pattern. Supplementary Figure S1 separates the left-and right-hemisphere results to show that the main conclusions do not depend on this averaging step. The qualitative pattern is similar across hemispheres, although the magnitudes of exact-match rates, mean shifts, and adjacent reorderings vary somewhat across developmental intervals.

**Figure S1:**
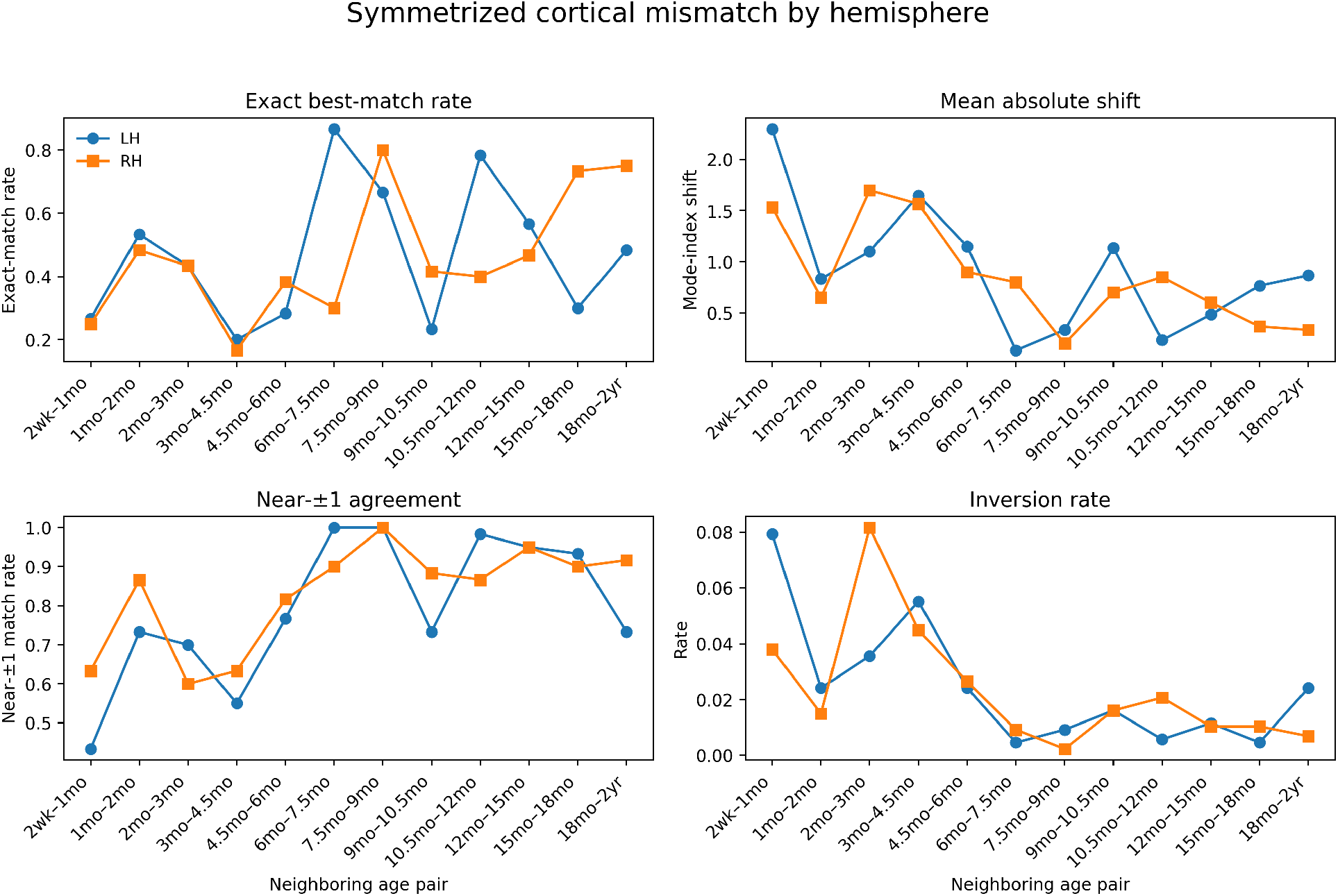
Symmetrized cortical mismatch summary by hemisphere. Directional neighboring-age mismatch metrics were symmetrized and then reported separately for the left and right hemispheres. The qualitative pattern is broadly similar across hemispheres, but the magnitude of exact-match rates, mean absolute shifts, near-neighbor agreement, and adjacent reorderings differs somewhat across developmental intervals. These hemisphere-specific results support the robustness of the main conclusion that independently computed age-specific LB modes are not directly comparable by index alone.

### S8 Supplementary view of local eigenvalue spacing (mode crowding)

In the main text, we summarize the relationship between local eigenvalue crowding and exact same-index matching after pooling across hemispheres and neighboring-age comparisons. Supplementary Figure S2 provides a more detailed view by showing hemisphere-specific crowding heatmaps and, in addition to exact matching, the corresponding near-neighbor match probabilities. This helps clarify that while exact same-index matching declines in crowded neighborhoods, much of the remaining agreement is still local.

**Figure S2:**
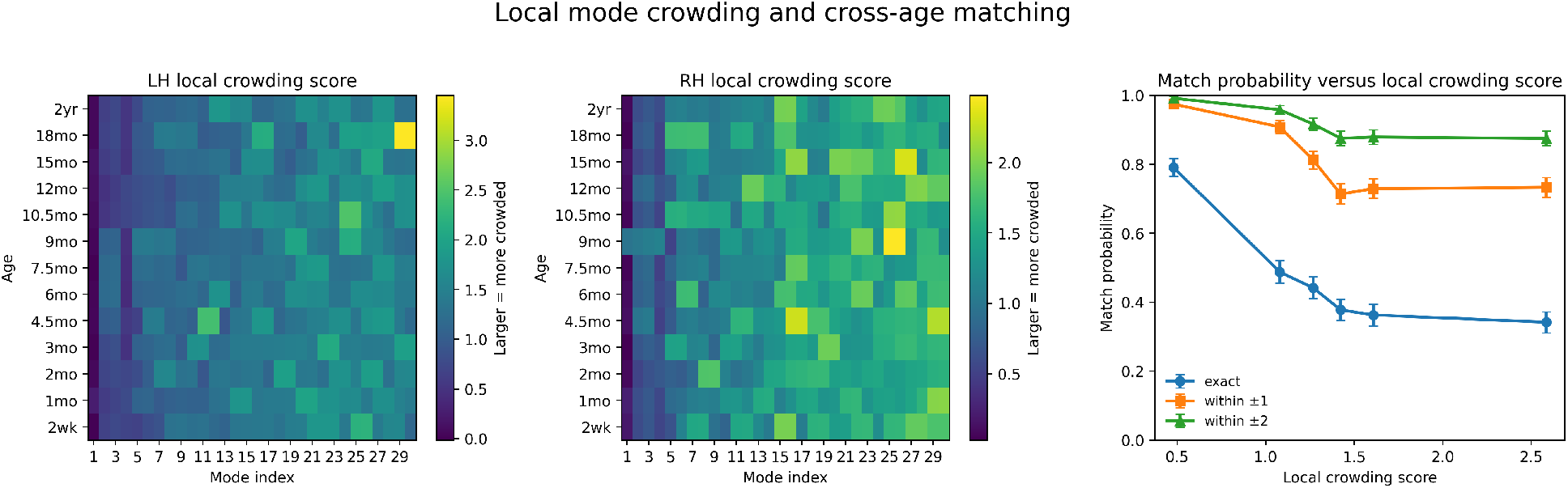
Supplementary view of mode crowding and cross-age matching. The left and middle panels show hemisphere-specific crowding-score heatmaps across age and mode index. The right panel shows the probability of exact matching and matching within *±*1 and *±*2 neighboring mode indices as functions of the crowding score. Although exact same-index matching declines most strongly in crowded neighborhoods, near-neighbor matching remains substantially higher, indicating that cross-age instability is often local rather than catastrophic.

### S9 Sequential Procrustes tracking improves same-index coefficient retention

As a downstream validation of the tracking analysis, we repeated the coefficient-space mismatch analysis using the Procrustes-tracked cortical basis and compared the results with those obtained from the raw, independently computed neighboring-age bases. The purpose of this supplementary analysis is not to introduce a new criterion for tracking, but to verify that the improved basis-level same-index correspondence documented in the main text also propagates to the coefficient level.

Supplementary Figure S3 shows this comparison for the most directly relevant coefficient-space summary: the fraction of squared coefficient mass remaining at the same nominal mode index after re-expression in a neighboring-age basis. We report this quantity for both *single-mode test patterns*, in which the source-age coefficient vector contains a single active mode, and *neighboring-mode test patterns*, in which the source-age coefficient vector is concentrated in a small neighborhood of nearby mode indices. In both cases, the plotted quantity is the exact same-index retained fraction before versus after Sequential Procrustes tracking.

The improvement is systematic. For the single-mode test patterns, exact same-index coefficient retention increased for all neighboring-age pairs after tracking, with especially large gains in intervals that had shown strong basis-level mismatch in the main analysis, including 2wk–1mo, 3mo–4.5mo, 9mo–10.5mo, and 12mo– 15mo. For the neighboring-mode test patterns, the same qualitative pattern held: exact same-index retention increased throughout, although the gains were somewhat smaller in magnitude, as expected for source patterns that already distribute coefficient mass across a small local neighborhood rather than a single exact mode.

These downstream results strengthen the interpretation of the main tracking analysis. Sequential Procrustes tracking does not merely improve overlap-based summaries of neighboring-age basis correspondence; it also improves the practical stability of neighboring-age coefficient representations. In that sense, the figure provides an important validation that the tracked basis is not only more coherent geometrically, but also more useful as a coordinate system for representing cortical patterns across age.

**Figure S3:**
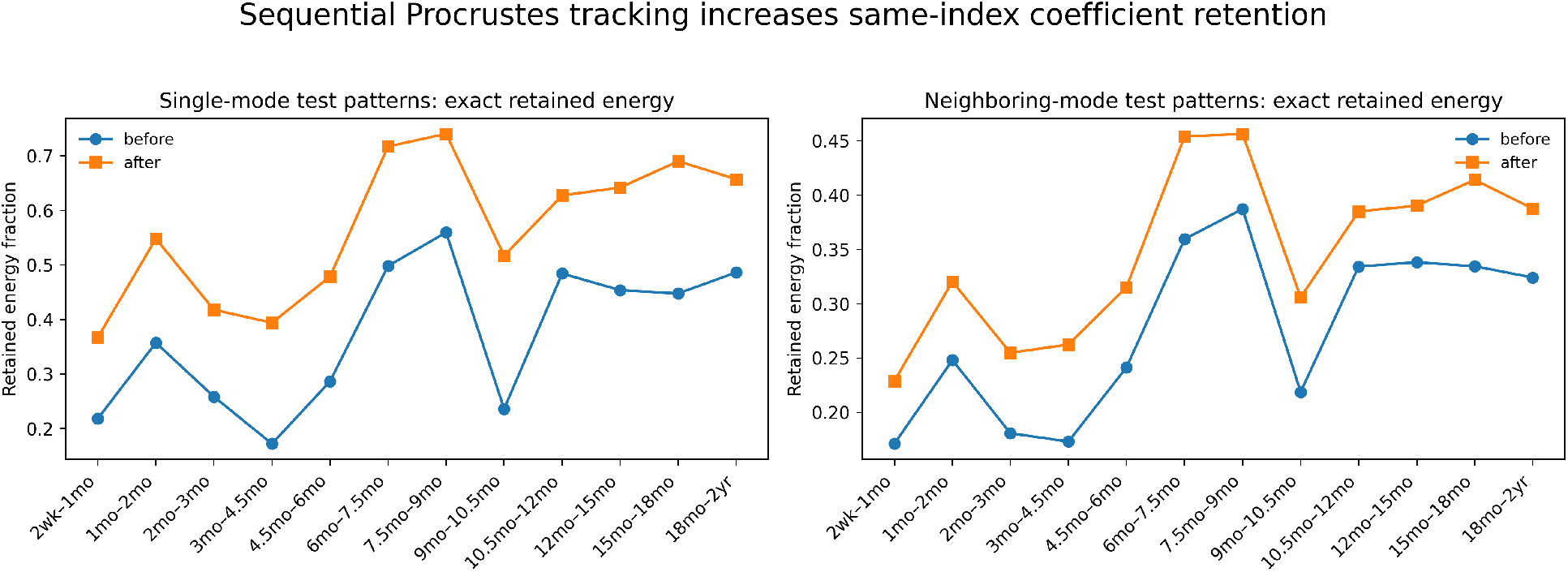
Sequential Procrustes tracking increases exact same-index coefficient retention. For each neighboring-age pair, curves compare the fraction of squared coefficient mass remaining at the same nominal mode index (i.e., “exact retained energy”) before and after Sequential Procrustes tracking. The left panel shows *single-mode test patterns*, in which the source-age coefficient vector contains a single active mode. The right panel shows *neighboring-mode test patterns*, in which the source-age coefficient vector is concentrated in a small neighborhood of nearby mode indices. In both pattern families, exact same-index coefficient retention increases systematically after tracking, indicating that the main effect of the procedure is to sharpen local neighboring-age correspondences into more exact same-index alignment. This figure serves as a downstream coefficient-level validation of the overlap-based tracking results reported in the main text.

